# Glu1, a conserved GH30_3 β-(1,6)-glucanase requires its catalytic residues for full virulence in *Ustilago maydis*

**DOI:** 10.64898/2026.10.29.762955

**Authors:** María Dolores Pejenaute-Ochoa, Ismael Moreno-Sánchez, Ramón Ramos Barrales, José Ignacio Ibeas

## Abstract

Plant-pathogenic fungi cause plant diseases and severe agricultural losses. The fungal cell wall is the first point of contact and defense barrier between fungus and host, maintaining cellular integrity and structural stability but also playing a dynamic role during infection. Through pathogenic development, the fungal cell wall must be continuously remodelled to accommodate morphological transitions and environmental challenges through diverse cell wall–modifying enzymes necessary for successful infection. Here, we characterise Glu1, a putative endo-β-(1,6)-glucanase (GH30) from the maize pathogen *Ustilago maydis*, conserved across Ustilaginales. Structural comparison with characterised GH30 crystal structures shows Glu1 most closely resembles GH30_3 β-(1,6)-glucanases. Glu1 expression is induced during infection, localises to the plasma membrane in close association with the fungal cell wall and is detected in the secreted fraction. Deletion of *glu1* reduces virulence, impairs plant colonization and tumour formation, and alters septation and cell morphology under saprophytic growth. Notably, genetic rescue with Glu1 catalytic-site variants largely restores saprophytic phenotypes but not virulence, suggesting a catalytic-dependent function of Glu1 specifically required for infection. Umag_04357, the other GH30 enzyme identified in *U. maydis*, reveals a conserved GH30 domain, including the catalytic residues, but is expressed at lower levels during infection and its deletion does not affect virulence. Together, our findings indicate that Glu1 is a conserved GH30 enzyme with a β-(1,6)-glucanase-like pocket whose catalytic residues are required for full virulence. To our knowledge, it is the first GH30 enzyme linked to virulence in Basidiomycota and could be considered a potential target for smut disease control.

## 1. INTRODUCTION

The smut fungi of the *Ustilaginaceae* family represent a diverse group of biotrophic pathogens that infect grasses (Xia et al., 2020). Among them, *Ustilago maydis* is a consolidated model system for studying plant–fungus biotrophic interactions in *Basidiomycota* (Dean et al., 2012). *U. maydis* is a dimorphic fungus which grows in its haploid phase as a saprophytic yeast while pathogenic development starts on the plant surface, where two compatible haploid cells recognise each other through pheromone-receptor signalling (Kämper et al., 2006). This interaction triggers the formation of conjugation tubes and subsequent cell fusion, resulting in a dikaryotic filament (Feldbrügge et al., 2004; Castanheira and Pérez-Martín, 2015). These filaments then differentiate into appressoria (Mendoza-Mendoza et al., 2009) and the fungus proliferates and colonizes plant tissue inducing tumour formation, the hallmark symptoms of corn smut. During the infection cycle, the fungal cell wall not only provides mechanical stability and protection from environmental stress, but also undergoes dynamic remodelling (Geoghegan et al., 2017; Gow and Lenardon, 2023). These modifications support morphogenetic transitions such as filaments formation, plant cuticle penetration, successful host tissues invasion (Mendgen et al., 1996; Wang and Lin, 2012; Pejenaute-Ochoa et al., 2024) and plant immune responses evasion (Ma et al., 2018; Rizzi et al., 2021).

The fungal cell wall is composed mainly of polysaccharides such as glucan and chitin and proteins where most of them are O-glycosylated (Latgé, 2007; Geoghegan et al., 2017; Gow and Lenardon, 2023). Glucans constitute approximately 50–60% of the dry weight of fungal cell wall and consist mainly of long linear chains of β-1,3-glucans (Bowman and Free, 2006; Garcia-Rubio et al., 2020), which has been the most studied (Martin et al., 2007; Reyre et al., 2022) due to its abundance and its role in pathogenesis as a PAMP (Wagener et al., 2020). However, other polysaccharides such as β-1,6-glucans are also critical components of the fungal cell wall, (Mouyna et al., 2000; Garcia-Rubio et al., 2020) and are found exclusively in the cell walls of fungi and oomycetes (Wawra et al., 2016), making them promising targets for the control of plant fungal pathogens. In this context, β-1,6-glucans are essential for cell wall integrity in *Magnaporthe oryzae* (Li et al., 2023). Their disruption causes swollen hyphae, defective appressoria and reduced adhesion in *Colletotrichum graminicola* (Oliveira-Garcia and Deising, 2016) and their reduction impairs hyphal formation and modulates host immune responses in *Candida albicans* (Umeyama et al., 2006,Bekirian et al., 2024).

As the fungal cell wall is a dynamic structure, fungi employ endo-β-(1,6)-glucanases (EC 3.2.1.75, pustulanases), which cleave internal β-1,6-glucan bonds, releasing oligosaccharides of glucose. According to the CAZy database (http://www.cazy.org/), almost all known endo-β-(1,6)-glucanases belong to glycoside hydrolase families 5 and 30 (GH5 and GH30). Several endo-β-(1,6)-glucanases have been characterised in Ascomycota, including those of *N. crassa* (Oyama et al., 2002) and *Aspergillus fumigatus* (Boisramé and Gaillardin, 2009), the three secreted β-1,6-glucanases from *Trichoderma* that contribute to mycoparasitism (Lora et al., 1995; De La Cruz and Llobell, 1999; Montero et al., 2005), GcnA from *Epichloë festucae*, required for maintaining a mutualistic relationship with its grass host (Bryant et al., 2007), and MoGlu16, required for cell wall remodelling during pathogenesis in *Magnaporthe oryzae* (Wang et al., 2024). In contrast, endo-β-(1,6)-glucanases remain largely unexplored in Basidiomycota, with only a few examples reported to date, such as LePus30A from *Lentinula edodes*, involved in post-harvest fruiting body autolysis (Konno and Sakamoto, 2011), and GH30A from *Coprinopsis cinerea*, linked to cell wall extension in stipes (Liu et al., 2020). However, no β-(1,6)-glucanases have been identified or characterised in Basidiomycota plant pathogens such as smut fungi. Here, we present Glu1, a putative endo-β-(1,6)-glucanase (GH30) of *Ustilago maydis*, conserved across Ustilaginales and required for successful pathogenic development. To our knowledge, Glu1 is the first GH30 enzyme in Basidiomycota whose catalytic residues are required for pathogenesis, and its active-site pocket strongly resembles those of GH30_3 β-(1,6)-glucanases, pointing to β-1,6-glucan modification as a virulence mechanism in smut fungi.

## 2. RESULTS

### 2.1. Glu1 has been identified as a putative endo-1,6-β-glucanase conserved in Ustilaginales and required for full virulence

We identified Umag_05223, hereafter referred to as Glu1, in a previous screening for *U. maydis* glycoproteins implicated in pathogenesis (Marín-Menguiano et al., 2019) where its deletion caused one of the strongest phenotypes. Glu1 was identified as a putative endo-1,6-β-glucanase based on BLAST and phylogenetic analyses. The protein contains a conserved GH30 glycoside hydrolase domain (aminoacids 100–358) associated with endo-1,6-β-glucanase activity. Glu1 also contains a predicted N-terminal signal peptide (aminoacids 1–24), a putative protease cleavage site at residue 468, and several predicted O- and N-glycosylation sites **(Figure 1a, upper panel, Table S1)**. AlphaFold predicts a classical TIM barrel (α/β barrel) fold for Glu1 (100–358) **(Figure 1a, upper panel)**. Closer inspection of the secondary structure revealed a complete (β/α)_8_ barrel in Glu1, whose eighth helix (α8, residues ∼528–540) is separated from β8 (residues ∼391–394) by an insertion of about 133 residues (∼395–527) that includes an intrinsically disordered region (IDR, residues ∼433–508) **(Figure 1a, lower panel)**. Phylogenetic analysis showed that Glu1 is specifically conserved within the Basidiomycota, predominantly among Ustilaginales **(Figure 1b and S1, Table S1)** and, to a lesser extent, in some Exobasidiomycetes. Although Glu1 does not have any paralogs, we identified another protein, Umag_04357, which shares a GH30 domain in *U. maydis*.

**FIGURE 1.**
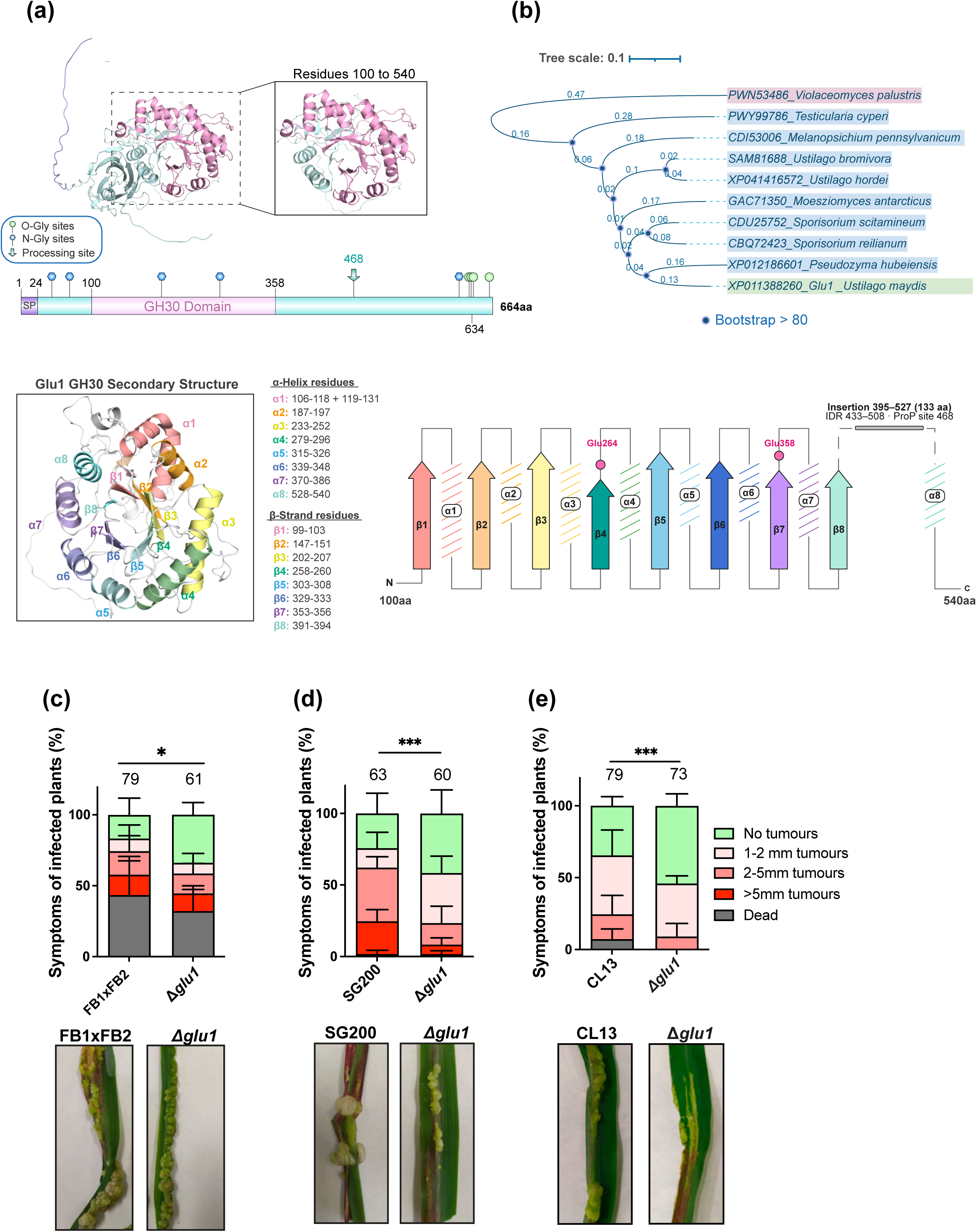
Glu1 has been identified as a putative endo-1,6-β-glucanase conserved in Ustilaginales and required for full virulence. **(a)** Upper panel: secondary structure of Glu1 TIM α/β barrel of the GH30 domain (upper) and domain architecture (lower): signal peptide (SP, 1–24, lilac), GH30 domain (100–358), predicted N- and O-glycosylation sites (blue and green lollipops), predicted processing site (residue 468, green arrow). Magnified GH30 extended domain (100-540aa). Lower panel: secondary structure of the GH30 domain, with α-helices α1–α8 and β-strands β1–β8 coloured salmon, orange, yellow, green, light blue, blue, purple and aquamarine and schematic representation of the GH30 domain, including the insertion between β8 and α8 and its associated IDR **(b)** Maximum-likelihood tree of Glu1 orthologues across Ustilaginales with *V. palustris* as outgroup. Scale bar, 0.1 substitutions per site; blue dots mark nodes with bootstrap >80 (1,000 replicates). **(c-e)** Symptoms of maize plants at 14 days post-infection (dpi) with Δ*glu1* mutants in FB1×FB2 **(c)**, SG200 **(d)** and CL13 **(e)**, compared with their wild types. Numbers above bars (plants scored); error bars, SD of three replicates; Mann–Whitney test (ns, not significant; *P < 0.05; ***P < 0.001).

To confirm the role of Glu1 in pathogenesis, we infected maize plants using Δ*glu1* deletion mutants in the sexually compatible *U. maydis strains* FB1 and FB2 (Herskowitz, 1989) **(Figure 1c)**, the solopathogenic strain SG200 (Bölker et al., 1995) **(Figure 1d)**, and the attenuated progenitor strain CL13 (Bölker et al., 1995) **(Figure 1e)**. Deletion of *glu1* impaired virulence in all backgrounds. As the effect was slightly more pronounced in CL13 **(Figure 1e)**, we reintroduced the wild-type (WT) *glu1* allele under its own promoter into the CL13 Δ*glu1*, which fully restored virulence, confirming the contribution of Glu1 to infection **(See Figure 6e)**.

### 2.2. Loss of Glu1 alters cell morphology and septation and impairs host colonization

To investigate the role of Glu1 in pathogenesis, we first examined whether deletion of *glu1* affects fungal morphology and/or cell division under saprophytic growth. In *U. maydis*, cytokinesis requires the sequential formation of a primary and secondary septum at the site of cell division, a process that depends on septin ring reorganization and coordinated deposition of chitin and glucans at the division site (Weinzierl et al., 2002). The Δ*glu1* mutant exhibited elongated cells and defects in septum formation during cell division **(Figure 2a, 2b and Table S2)**. Despite these abnormalities, no differences in growth rate were observed between the mutant and WT strains **(Figure S2a)**, indicating that cell cycle progression is not severely affected. These results suggest that Glu1 may contribute to cell wall remodelling during cytokinesis.

**FIGURE 2.**
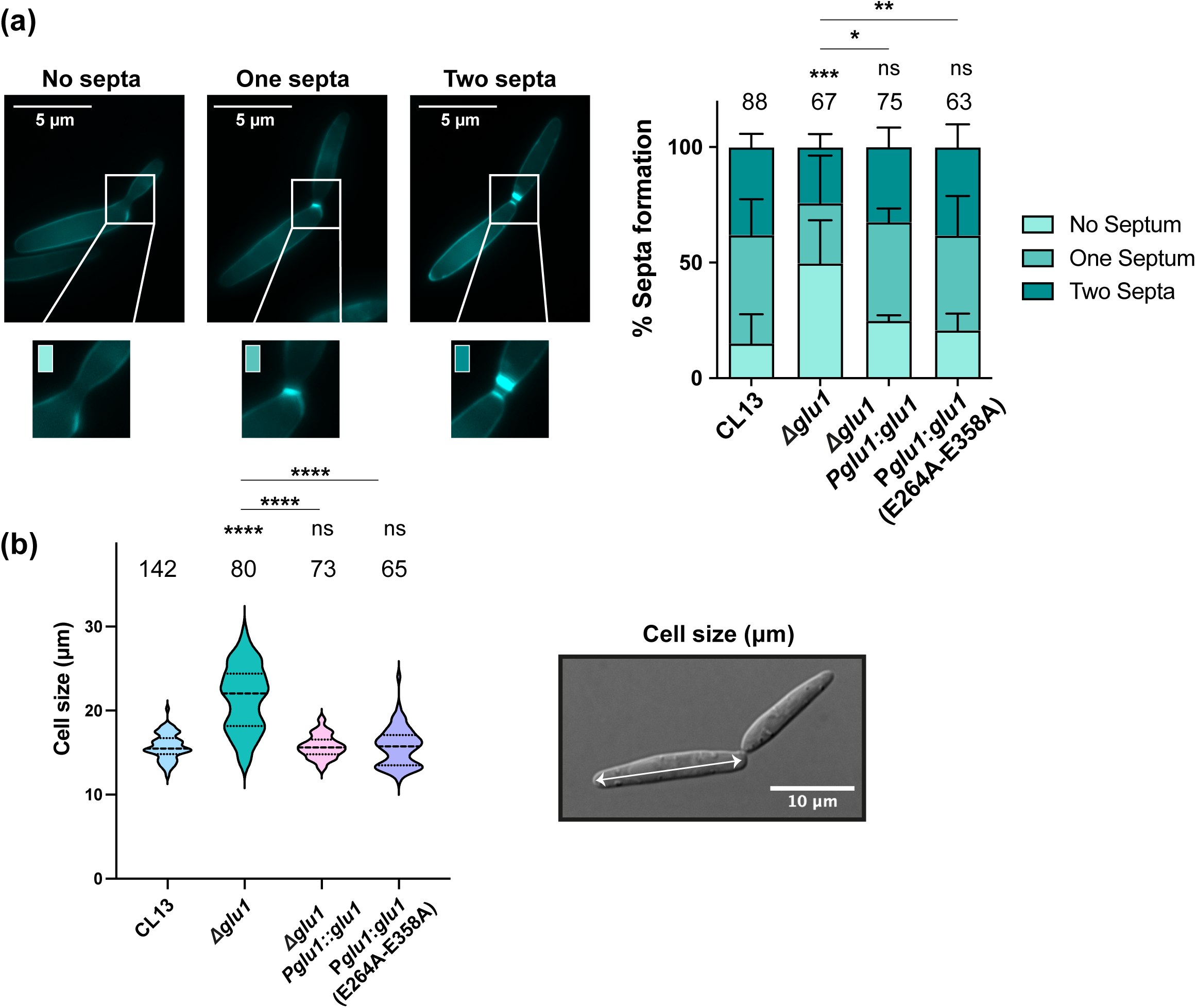
Loss of Glu1 alters septation and cell size, but these defects are not dependent on its catalytic residues. **(a)** Quantification of septum number (no septum, one septum, two septa) in CL13, Δ*glu1*, Δ*glu1* P*glu1*:*glu1*, and Δ*glu1* P*glu1*:*glu1*(E264A/E358A) strains grown under saprophytic conditions and stained with Calcofluor White (CFW). Statistics were performed using one-way ANOVA followed by Dunn’s post-hoc test for multiple comparisons. **(b)** Quantification of cell size (μm) at the point of division for the same strains. Numbers above bars (cells scored). Error bars, SD from at least two independent replicates. Mann–Whitney test (ns, not significant; *, P < 0.05; **, P < 0.005; ***, P < 0.0005; ****, P < 0.0001).

We next analysed which stages of fungal pathogenesis could be affected. Expression of *glu1* is induced during infection starting at 1 day post-inoculation (dpi), reaching maximum expression between 3 and 5 dpi **(Figure 3a)**, corresponding to the plant colonization stage. In the FB1×FB2 background conjugation tube and mating formation were unaffected in Δ*glu1* mutants **(Figure 3b and 3c)**, suggesting that Glu1 is not required for sexual recognition or cell fusion. Given that Glu1 is dispensable for mating and its deletion leads to a more pronounced virulence defect in the CL13 **(Figure 1e)**, we focused on this genetic background, as its attenuated virulence facilitates the detection of modest differences in plant infection (Di Stasio *et al*., 2009; Djamei *et al*., 2011). Filamentation assays on PD-charcoal plates revealed that Δ*glu1* colonies were slightly whiter than WT, a phenotype restored to WT levels in the complemented strain **(Figure 3d)**. However, no defects were observed in appressorium formation on plant **(Figure 3e)**. These results indicate that early developmental steps required for host penetration are not compromised in the Δ*glu1* mutant. Therefore, we analysed fungal proliferation by measuring fungal biomass at 3 dpi. Δ*glu1* showed a reduction in mean fungal biomass compared to the WT strain during the colonization step, although this difference was not statistically significant (*P* = 0.1), possibly because this approach lacks the sensitivity to detect subtle phenotypes **(Figure S2b)**. Consistently with this trend, microscopic analysis revealed altered fungal morphology in infected plant tissue. While wild-type hyphae extended longitudinally along plant cells forming interconnected networks, the Δ*glu1* mutant showed compact, disorganised hyphal aggregates with limited tissue invasion **(Figure 3f left)**. Quantification of WGA-AF488-stained fungal signal at 3 dpi showed a lower colonization in Δ*glu1* than in the WT strain **(Figure 3f right)**. This aberrant hyphal architecture may reflect defects in cell wall remodelling during invasive growth, according to previous length and septation phenotypes **(Figure 2a and 2b)**.

**FIGURE 3.**
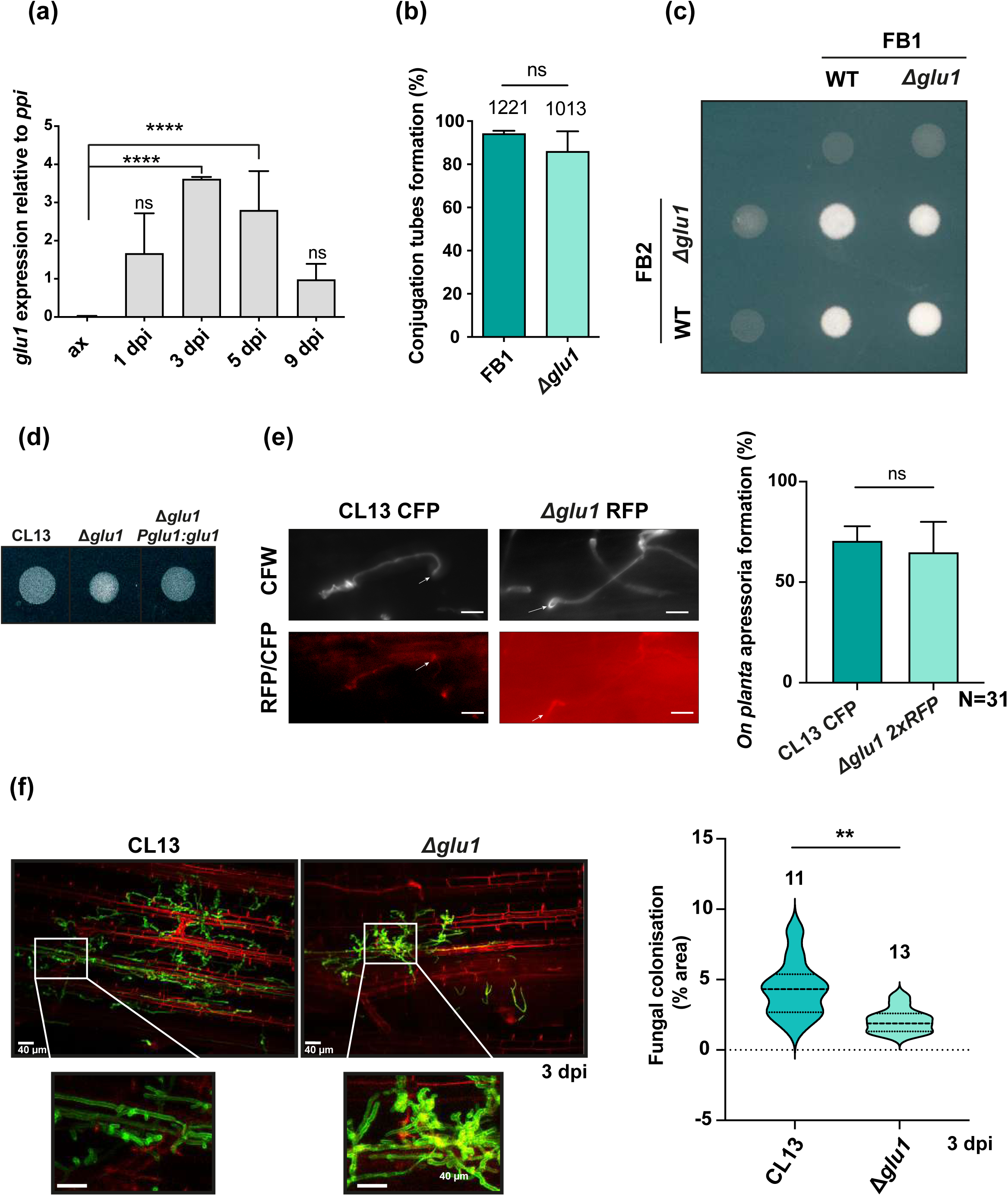
Glu1 is dispensable for mating and appressorium formation but required for efficient plant colonization. **(a)** *glu1* expression relative to *ppi1* (qRT-PCR) in axenic culture (ax) and in maize at 1, 3, 5 and 9 days post-infection (dpi). **(b)** Conjugation tube formation (%) in FB1 and Δ*glu1* upon pheromone induction. **(c)** Mating assay on charcoal plates (fuzzy colonies) of crosses between FB1 and FB2 wild-type and Δ*glu1* strains. **(d)** Filamentation of CL13 and Δ*glu1* on PD-charcoal plates. **(e)** *In planta* appressoria of CL13-CFP and Δ*glu1*-2xRFP stained with Calcofluor White (CFW) (arrows); quantification of appressorium formation (%) on the right (N = 31). **(f)** Representative confocal images of hyphal colonization at 3 dpi (right) and quantification of fungal plant colonization. Numbers above bars (images scored); Error bars, SD from two **(b,e,f)** or three **(a)** independent replicates. Student’s t-test: ns, not significant; *P < 0.05; **P < 0.005; ***P < 0.0005; ****P < 0.0001.

### 2.3. Glu1 is a secreted protein that localises to the plasma membrane in close association with the cell wall

To better understand the role of Glu1 in cell wall dynamics and host colonization, we investigated its subcellular localisation. Since expression of *glu1* under its native promoter resulted in weak fluorescence signals, likely due to its low expression levels during saprophytic growth **(Figure S3)**, we expressed a GFP-tagged version of Glu1 under the constitutive *otef* promoter. Fluorescence microscopy revealed that Glu1-GFP localised to the cell periphery. Co-staining with Calcofluor White (CFW), which labels chitin and β-glucans in the fungal cell wall, showed a close spatial association between Glu1 and the cell wall **(Figure 4a, upper panel)**. Interestingly, in protoplasts, where the cell wall was enzymatically removed, Glu1-GFP remained associated with the plasma membrane **(Figure 4a, lower panel)**, suggesting that Glu1 localises to the membrane and is positioned at the interface with the cell wall, consistent with its predicted function as a β-(1,6)-endoglucanase. As Glu1 is predicted to contain a putative protease cleavage site and a signal peptide **(Figure 1a)**, we investigated its processing and secretion. Western blot analysis of total protein extracts revealed two bands: a major band of ∼100 kDa corresponding to the full-length Glu1-GFP fusion, and a fainter band of ∼50 kDa, consistent with the putative processed form of the protein retaining the GFP tag **(Figure 4b)**. This putative processed site lies in the long segment between β8 and α8 of the GH30 barrel **(Figure 1a**, **Figure 4b)**, and it is not predicted in the closest orthologues. It is therefore unclear whether this band results from specific processing or from protein breakdown. To assess whether Glu1 is also secreted, we performed a colony secretion assay, detecting secreted proteins diffusing into the underlying agar; Glu1-GFP signal was detected in the extracellular fraction **(Figure 4c)**, supporting that Glu1 is secreted.

**FIGURE 4.**
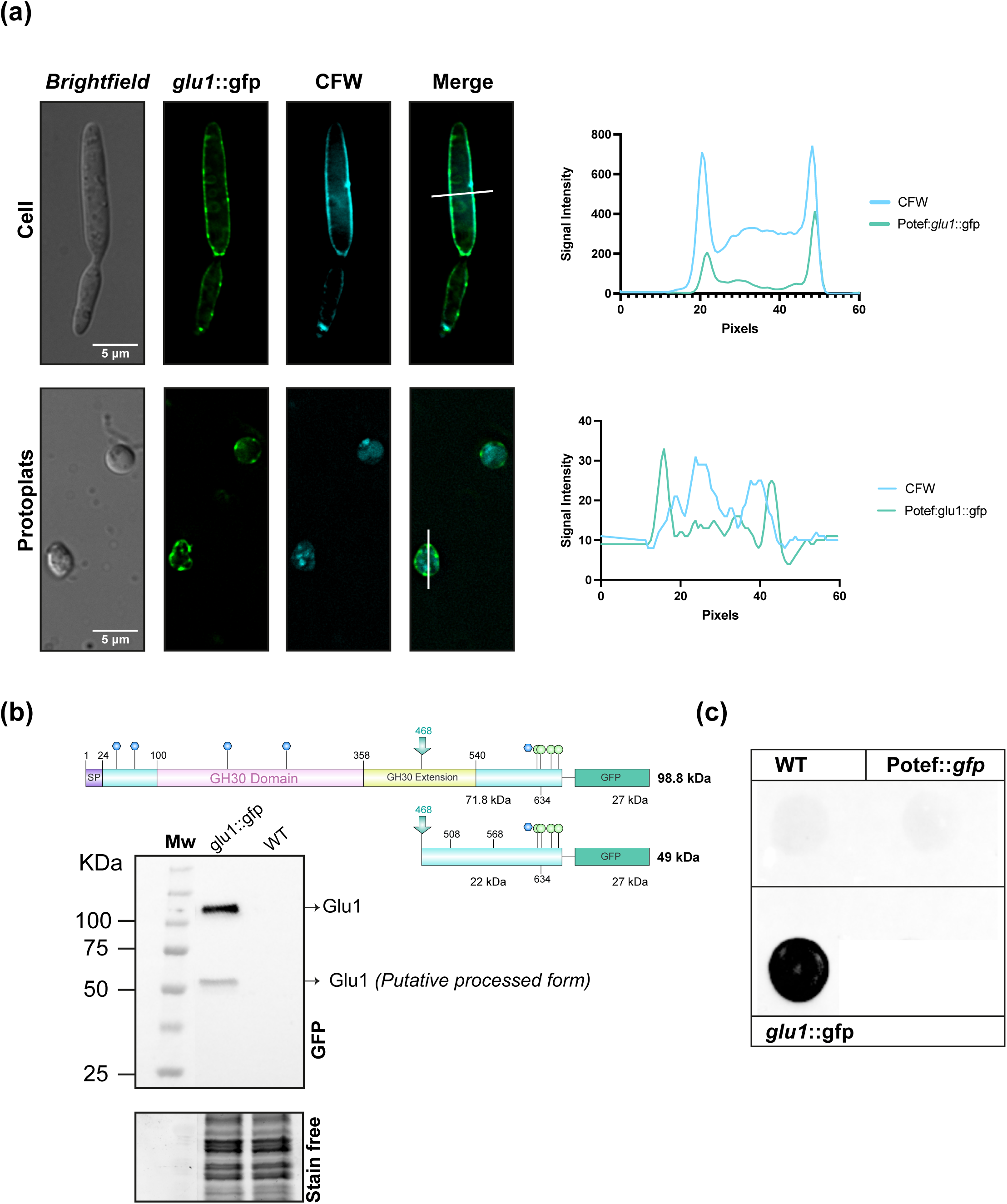
Glu1 localises to the plasma membrane and is secreted. **(a)** Subcellular localisation of Glu1::GFP expressed under the constitutive P*otef* promoter in cells (top) and protoplasts (bottom), co-stained with CFW (scale bars, 5 μm). Right: GFP and CFW intensity profiles along the white line in the merged images, for a representative cell (top) and protoplast (bottom). **(b)** Schematic representation of Glu1::GFP protein with the predicted sizes of the full-length (71.8 kDa Glu1 + 27 kDa GFP) and putative processed (22 kDa C-terminal fragment + 27 kDa GFP) forms, and corresponding anti-GFP western blot of total protein from *glu1*::*gfp* and WT strains under saprophytic conditions, showing the full-length fusion (∼100 kDa), and the ∼50 kDa form. Stain-free staining was used as loading control. **(c)** Colony secretion assay of Glu1::GFP on PD-charcoal plates under pathogenic conditions; SG200 WT and SG200 expressing cytoplasmic GFP (P*otef*) are cell-lysis controls.

### 2.4. The (β/α)₈ GH30 fold and the predicted catalytic-sites residues of Glu1 are conserved across Ustilaginales

Given its localisation in close proximity to the cell wall, consistent with a role in cell wall modification, we aimed to further investigate its catalytic domain and substrate-binding pocket through structural analysis.

Using AlphaFold we identified proteins structurally similar to Glu1 that share the conserved GH30 glycoside hydrolase domain. Among these, BT3312 from *Bacteroides thetaiotaomicron*, a GH30_3 β-(1,6)-endoglucanase that has been crystallised and functionally characterised (Temple et al., 2017) showed the highest structural similarity to Glu1 (E-value = 8.87 × 10⁻³⁵; 20.3% sequence identity). BT3312 revealed a TIM barrel catalytic domain with a deep substrate-binding pocket adapted to interact with 1,6-β-glucan, where its catalytic residues, BT-Glu238 and BT-Glu339, act as the general acid/base and nucleophile, respectively, of the clan GH-A catalytic mechanism (Kumar et al., 2019). Structural alignment of the full-length Glu1 (mean pLDDT 81.25, Figure S4) and BT3312 showed a root-mean-square deviation (RMSD) of 1.28 Å. The first seven α-helices and eight β-strands of Glu1, which form the (β/α) barrel, aligned with BT3312. However, this value was influenced by regions of Glu1 that have no counterpart in BT3312, including a long internal segment (residues ∼395–527) and the N- and C-terminal extensions. Closer inspection of the secondary structure then showed that the eighth α-helix (α8) of Glu1, although located after this long insertion (residues ∼528–540), occupied the position of α8 in BT3312. Consistently, extending the core to include it (residues 100–540) maintained a low RMSD (1.07 Å), confirming a complete (β/α)₈ barrel in Glu1 **(Figure 1a lower panel, Figure 5a and Table S3)**.

**FIGURE 5.**
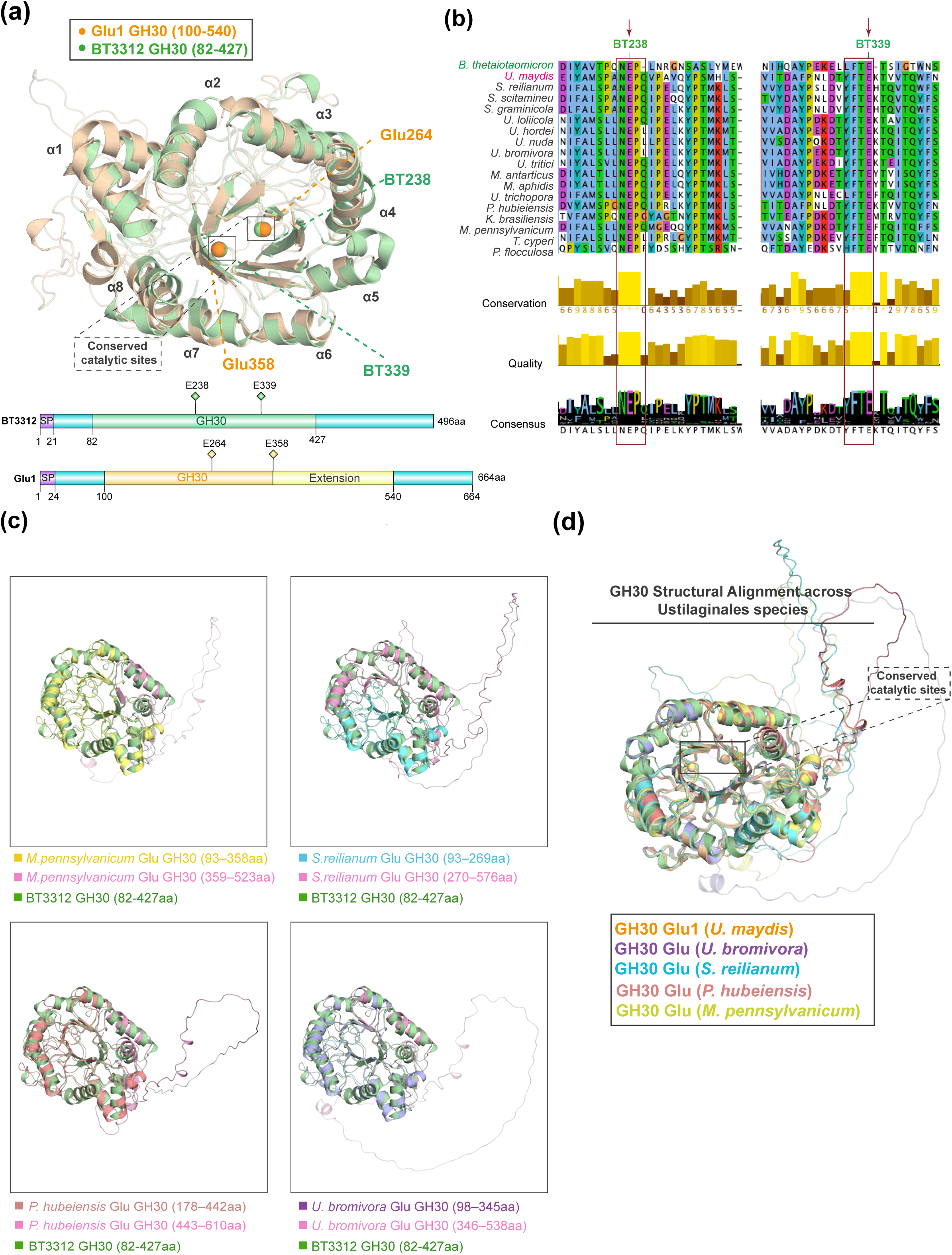
The extended GH30 domain and catalytic residues of Glu1 are conserved across Ustilaginales and in BT3312. **(a)** Upper panel: structural alignment of the Glu1 GH30 domain (wheat, 100–540) and the crystal structure 5NGL of *B. thetaiotaomicron* BT3312 (green, 82–427), highlighting the catalytic residues BT-Glu238 and BT-Glu339 of Glu1, equivalent to Glu264 and Glu358 (spheres), and α-helices α1–α8. Lower panel: domain architecture of BT3312 (496aa, predicted: GH30 82–427) and Glu1 (664a, predicted: GH30 100–358, extended to 540), with the signal peptide (SP, purple) and the catalytic residues (diamonds). **(b)** Multiple sequence alignment of the GH30 domain of BT3312 across Ustilaginales, showing the catalytic residues (arrows, boxes). Conservation, quality, and consensus tracks are shown below the alignment. **(c)** Structural alignment of the GH30 domain across Ustilaginales: *M. pennsylvanicum* (yellow, 93–358), *S. reilianum* (cyan, 93–269), *P. hubeiensis* (salmon, 178–442) and *U. bromivora* (violet, 98–345) with BT3312 (green, 82–427); the structurally extended region of each orthologue is in pink. **(d)** Superposition of the GH30 domains of Glu1 and the Ustilaginales species showed in c).

Sequence and structural alignment showed that the overall architecture is conserved across Ustilaginales, with Glu264 and Glu358 occupying the same position as BT3312’s BT-Glu238 and BT-Glu339 in all orthologues **(Figure 5b and c, Figure S1, S5 and Table S3)**. While the equivalent of Glu264 always falls within the GH30 domain annotated by InterPro, the equivalent of Glu358 lies outside it in some orthologues **(Figure S1)**. Structural alignment of BT3312 with some of the most representative orthologues (*S. reilianum*, *U. bromivora*, *P. hubeiensis* and *M. pennsylvanicum*) showed that the GH30 domain is longer than annotated. The downstream region contributes the seventh α-helix and the eighth β-strand and α-helix (α8), which lie beyond an insertion of 122–176 residues absent from BT3312 placing the second catalytic residue within the structure in the seventh β-strand **(Figure 5c and d, Table S3)**. In some orthologues, secondary structure assignment did not detect all the expected elements. However, analysis of the residues at the equivalent position in BT3312, together with their Cα geometry and high pLDDT (Figure S6), showed canonical α-helices and β-strands, indicating a limitation of the structure assignment rather than a loss of fold **(Table S3)**. Together, the conserved catalytic glutamates and the similarity of the GH30 domain of Glu1 and its Ustilaginales orthologues to BT3312 identify Glu264 and Glu358 as the catalytic acid/base and nucleophile of Glu1 suggesting a β-1,6-glucanase activity.

### 2.5. Glu1 has a β-1,6-glucanase-type pocket whose catalytic residues are required for infection

To test whether the Glu1 pocket can accommodate β-1,6-glucan in proximity to the catalytic residues, we performed molecular docking with β-gentiobiose, a β-1,6-linked glucose dimer that mimics the natural substrate of β-(1,6)-endoglucanases (Plakys et al., 2024). Docking analysis revealed that β-gentiobiose was accommodated within a well-defined binding pocket of the TIM barrel fold, formed by nine residues within 4.5 Å of the ligand: Asp156, Glu264, Tyr272, Tyr335, Glu358, Thr360, Val361, Val362 and Trp395 **(Figure 6a, Table S4)**. Five of them (Glu264, Glu358, Tyr335, Thr360 and Trp395) made polar contacts with the ligand (≤3.2 Å). The two following poses placed the ligand in the same pocket (centroid displacement ≤1.2 Å) with the same residues within 4.5 Å, but in a different orientation (RMSD 2.9–3.5 Å). Polar contacts with Glu264 and Glu358 were conserved in all three poses, at distances of 2.7–2.9 Å, a positioning compatible with catalysis **(Figure 6a and Table S4)**. The predicted binding free energy (ΔG) was −6.2 kcal/mol, similar to that obtained for the BT3312–gentiobiose docking (−6.8 kcal/mol) **(Table S5),** indicating that Glu1 pocket can accommodate β-1,6-glucan next to the catalytic residues, as in BT3312.

**FIGURE 6.**
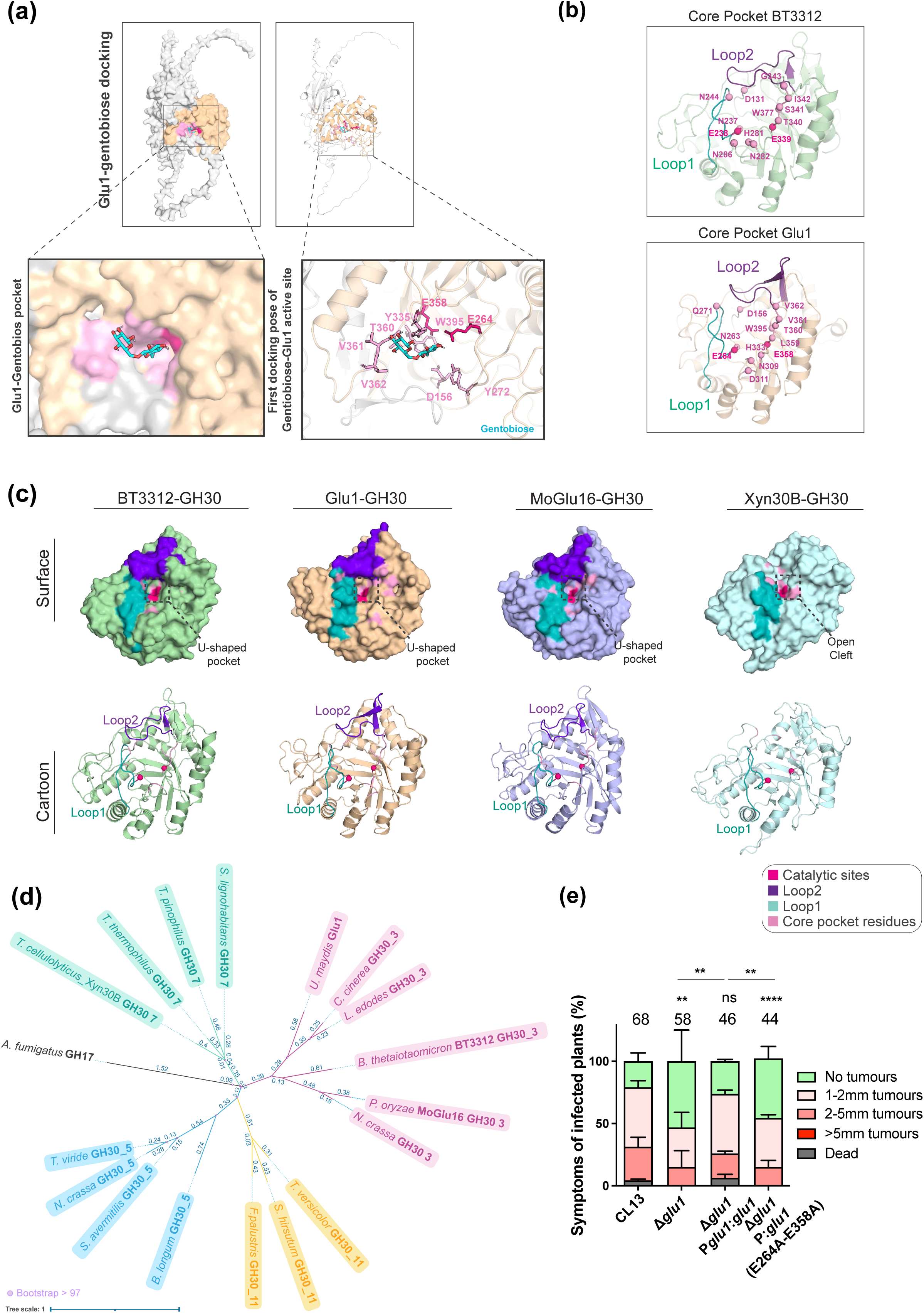
Glu1 docks β-1,6-gentiobiose in a β-1,6-glucanase-like pocket, and its catalytic residues are required for full virulence. **(a)** Docking of β-gentiobiose (cyan sticks) onto Glu1: surface (left) and cartoon (right) views, with the GH30 domain in wheat, pocket residues in pink and catalytic residues Glu264/Glu358 in hot pink; boxed regions are magnified below (pocket residues within 4.5 Å of the first pose are labelled). **(b**) Core pocket of BT3312 (green) and Glu1 (wheat), with Loop 1 (teal) and Loop 2 (violet); labels give BT3312/Glu1 residue pairs. **(c)** Surface (top) and cartoon (bottom) of the GH30 domain of BT3312, Glu1, MoGlu16 and Xyn30B, with catalytic sites (magenta), Loop 1 (teal), Loop 2 (purple) and core pocket residues (pink). BT3312, Glu1 and MoGlu16 have a U-shaped pocket, whereas Xyn30B has an open cleft. **(d)** Maximum-likelihood phylogeny of characterised fungal and bacterial members of GH30_3 (pink), GH30_5 (blue), GH30_7 (green) and GH30_11 (yellow), including Glu1 and BT3312. Scale bar, 0.1 substitutions per site; violet dots mark nodes with bootstrap support >97 (1,000 replicates). A GH17 enzyme were used as an outgroup **(e)** Symptoms of maize plants infected with CL13, Δ*glu1*, Δ*glu1* P*glu1:glu1* and Δ*glu1* P*glu1:glu1*(E264A_E358A) at 14 dpi. Numbers above the bars (plants scored); error bars, SD of three replicates. Mann–Whitney test (ns, not significant; **P < 0.005; ****P < 0.0001).

To further assess whether the Glu1 pocket shares the structural features that give BT3312 its specificity for β-1,6-glucan, we compared both structures **(Figure 6b and Table S6)**. As BT3312 is a bacterial enzyme, we also included MoGlu16, a fungal GH30_3 from *Magnaporthe oryzae* (Wang et al., 2024) to test whether these features are shared by fungal homologues. In BT3312, two loops close the active-site cleft, leaving room only for the U-shaped, hooked conformation of β-1,6-glucan and excluding the more linear glucans and xylan (Temple et al., 2017). Both loops aligned closely with equivalent regions in Glu1 and MoGlu16. Loop 1 (BT3312 236–252) corresponded to Glu1 262–279 and to MoGlu16 234–251, and loop 2 (BT3312 381–398), rigidified by the Cys393–Cys396 disulfide, to Glu1 399–418 and MoGlu16 367–386, which contain a matching disulfide (Cys415–Cys418, 2.02 Å, and Cys381–Cys384, 2.01 Å, respectively) **(Figure 6b and Table S6)**. The catalytic glutamates, the −1 subsite residues (Trp179, Asn237 and Trp377) and the positive-subsite Asn282 were also conserved in both proteins **(Figure 6b and Table S6)**. Of the two residues that distinguish BT3312 from other GH30 subfamilies, Asp131 was conserved (Asp156 in Glu1, Asp128 in MoGlu16), whereas Asn244 had no equivalent and only chemically similar candidates further away (Gln271 and Asn241, both 4.3 Å) **(Table S6)**. These comparisons show that Glu1 conserves the catalytic glutamates, both loops (including the loop 2 disulfide), the diagnostic Asp (Asp156 in Glu1), the −1 subsite residues and the catalytic glutamates. Only a few positive-subsite residues are replaced or displaced, supporting a pocket organised like that of BT3312.

Besides GH30_3, which includes β-1,6-glucanases such as BT3312, fungal GH30 proteins fall into three further subfamilies: GH30_7 (xylanases and glucuronoxylanases), the closest to GH30_3, and the more distant GH30_5 and GH30_11 (β-1,6-galactanases) (Li et al., 2022; St John et al., 2010). Among the structural homologues of Glu1 in the AlphaFold database, we found the GH30_7 xylanase Xyn30B from *Talaromyces cellulolyticus* (Nakamichi et al., 2019). To rule out a xylanase-type function, we compared Glu1 with Xyn30B. The catalytic domain of Glu1 is structurally closer to BT3312 than to Xyn30B: 236 of its 321 residues superimpose on BT3312 (RMSD 1.07 Å), but only 110 on Xyn30B (RMSD 1.46 Å) **(Table S6, Figure 6c)**. The two active sites differ in shape: in BT3312 and Glu1 the catalytic site lies in a U-shaped pocket enclosed by two loops, whereas in Xyn30B loop 1 is present but loop 2 has no equivalent, leaving a more open cleft that accommodates the linear xylan chain **(Figure 6c)**. The catalytic glutamates (Glu202 and Glu297) and the −1/+1 subsite residues Trp141, Asn201 and Tyr279 of Xyn30B matched identical residues in Glu1 (Glu264, Glu358, Trp208, Asn263 and Tyr335), MoGlu16 and BT3312 **(Table S6)**. As these positions are conserved across different GH30 subfamilies, they do not discriminate between activities. The fourth subsite residue, Tyr209, matched Tyr272 in Glu1, whereas BT3312 and MoGlu16 have Ser245 and Met244 at this position. In contrast, the elements that define xylanase specificity were absent from Glu1 and MoGlu16. The positions equivalent to Arg46, which recognises the MeGlcA substituent, and Asn93, associated with xylobiohydrolase activity, were occupied by different residues in Glu1 (Val108 and Gly158 in Glu1); most subsite −2b residues were not conserved and the Cys241–Cys242 disulfide that rigidifies the xylanase-specific loop was replaced by Ala305 and Trp306 **(Table S6)**. Together, these comparisons argue against a GH30_7 xylanase function and support β-1,6-glucanase (GH30_3) as the activity that best matches the Glu1 pocket. Consistently, a phylogenetic analysis of representative GH30_3, GH30_5, GH30_7 and GH30_11 members placed Glu1 within the GH30_3 clade **(Figure 6d)**.

Consistent with these structural predictions, molecular docking with other potential substrates β-1,4-xylan (xylanase activity), β-1,3-glucan (alternative glucanase activity), and chitin-derived oligomers (chitinase activity) showed lower binding affinities **(Table S5)**. Together with the conservation of the catalytic glutamates across Ustilaginales and the similarity of the pocket to that of BT3312, these results support Glu264 and Glu358 as the catalytic residues of Glu1 and β-1,6-glucan as its most likely substrate.

Having identified Glu264 and Glu358 as the catalytic residues, we asked whether they are required for infection. We substituted them with alanine (E264A and E358A) and introduced the mutated alleles into the CL13 Δ*glu1* mutant under Glu1 promoter. Infection assays in maize revealed that reintroducing the WT *glu1* gene restored the infective phenotype, while strains carrying the E264A and E358A substitutions showed infection defects similar to the Δ*glu1* mutant **(Figure 6e and Table S2)**.

These results indicate that Glu1 catalytic residues are required for infection. In contrast, the variants largely rescued cell size (median values approaching WT, although a subset of cells remained enlarged) and septation **(Figure 2a and 2b)**. This could reflect a non-catalytic role of Glu1 in cell morphology and septation, or residual activity of the variants that is sufficient under saprophytic growth, where *glu1* expression is low, but not during infection, when *glu1* is strongly induced **(Figure 3a)**.

### 2.6. Umag_04357, the second GH30 protein of *U. maydis*, is dispensable for virulence

Given the role of Glu1 in virulence, we next asked whether the other GH30 enzyme in *U. maydis* contributes similarly. We analysed the sequence and structure of Umag_04357. Umag_04357 contains a conserved GH30 domain (aminoacids 150–465), structurally similar to Glu1 and retains the key catalytic residues (Umag_04357-Glu317 and Umag_04357-Glu413) **(Figure 7a upper panel and Table S1)**. Despite these similarities, several features distinguish Umag_04357 from Glu1. Notably, Umag_04357 lacks a signal peptide and the C-terminal O-glycosylation motifs found in Glu1, has a predicted proteolytic cleavage site upstream of the GH30 domain (position 74) and is smaller in size **(Figure 7a lower panel)**. Structural alignment of the AlphaFold models of Glu1 and Umag_04357 showed that the GH30 (β/α)₈ barrels of Glu1 and Umag_04357 are well conserved (RMSD 0.71 Å over 295/350 core residues), whereas the full-length proteins differ substantially, (RMSD 0.77 Å, 316/602 residues superimposing) **(Figure 7a and Table S7)**. The eight β-strands and eight α-helices of the barrel are equivalent in both proteins, and the main difference within the barrel is the segment between β8 and α8, which is much shorter in Umag_04357 than in Glu1 (41 vs 133 residues) **(Table S7)**. Phylogenetic analysis revealed that Umag_04357 is conserved across Ustilaginales **(Figure 7b, and S7)**, and sequence alignments confirmed that the catalytic Glu1 residues are well preserved in these orthologues **(Figure 7c and S7)**. The two proteins cluster closely within the Ustilaginales clade (evolutionary distances of 0.54–0.82, versus 1.73 for the outgroup), although Glu1 appears slightly more divergent **(Figure S8)**. Both genes are expressed during colonization: *glu1* peaks at 2 dpi and *Umag_04357* at 4 dpi, although *Umag_04357* expression is substantially lower than that of *glu1* (Lanver et al., 2018) **(Figure 7d)**. To investigate whether Umag_04357 plays a role in virulence, we generated a deletion in CL13 and performed maize infection assays. The Δumag_04357 mutant showed no detectable differences in virulence compared to the WT strain **(Figure 7e)**. Together, these results indicate that, under the tested conditions, Glu1 and not Umag_04357 is the GH30 enzyme functionally coupled to virulence in *U. maydis*, despite their shared catalytic architecture.

**FIGURE 7.**
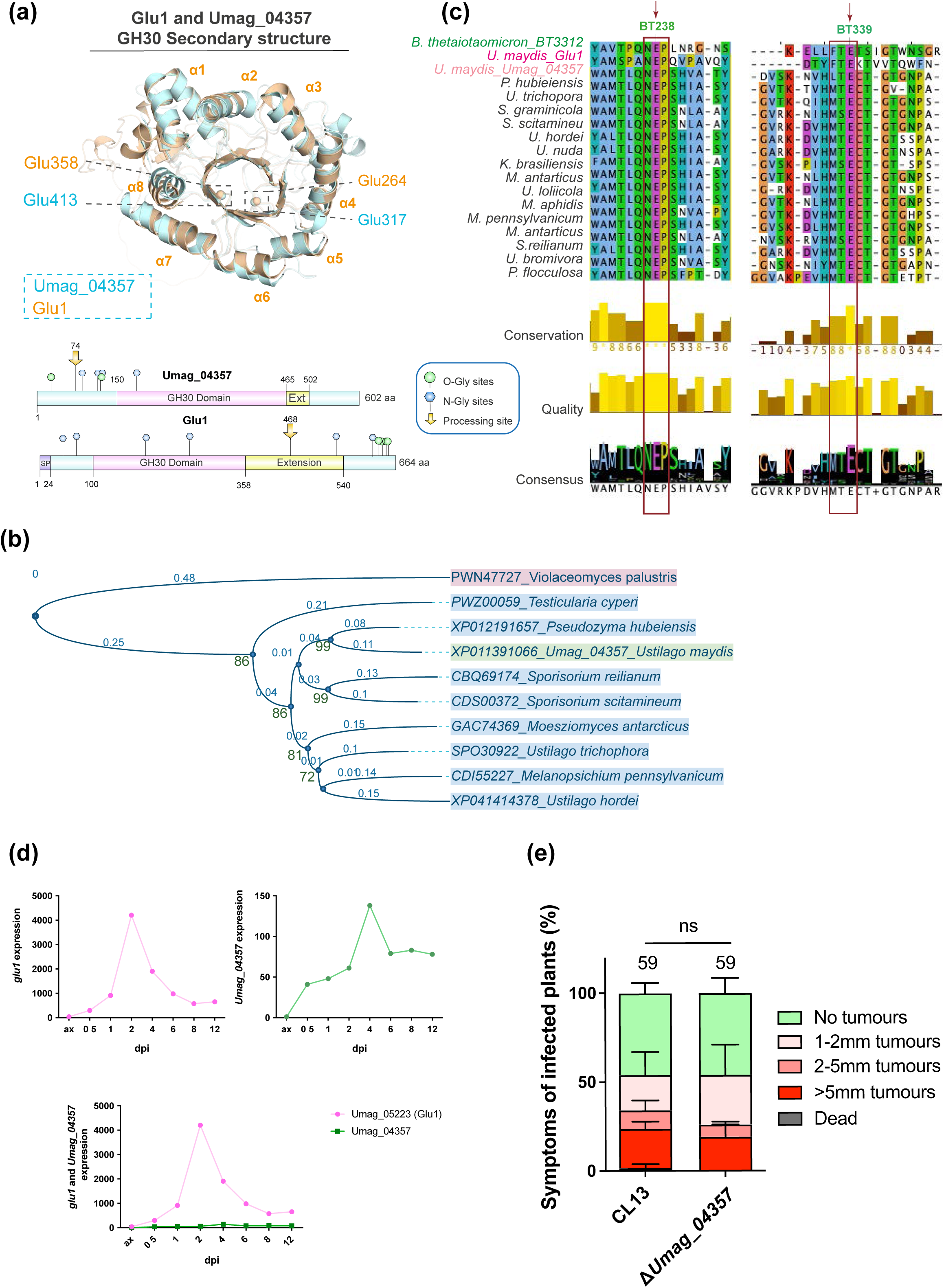
Umag_04357 shares structural and catalytic conservation with Glu1 but is dispensable for virulence. **(a)** Upper panel: Superposition of the extended GH30 domains of Glu1 (wheat) and Umag_04357 (cyan), showing the equivalent α-helices (α1–α8) and the conserved catalytic residues Glu264/Glu358 (Glu1) and Glu317/Glu413 (Umag_04357). Lower panel: domain architecture of Umag_04357 (602 aa, predicted GH30:150-465 extended to 502) and Glu1 (664 aa, predicted GH30: 100-358 extended to 540), with the GH30 domain, signal peptide, predicted N- and O-glycosylation sites and processing sites. **(b)** Multiple sequence alignment of the GH30 domain across Ustilaginales, showing conservation of the catalytic residues (magenta arrows/boxes) and their sequence context. Conservation, quality, and consensus tracks are shown below the alignment. **(c)** Maximum-likelihood phylogenetic tree showing conservation of Umag_04357 orthologues across Ustilaginales and related Basidiomycota, with *Violaceomyces palustris* as an outgroup. Scale bar, 0.1 substitutions per site; blue dots mark nodes with bootstrap >80 (1,000 replicates) marked with a blue dot. **(d)** Expression profiles of *glu1* (pink) and *Umag_04357* (green) during *U. maydis* growth in axenic culture (ax) and at different time points of maize colonization (0.5–12 dpi). **(e)** Quantification of symptoms in maize plants infected with CL13 and Δ*Umag_04357* strains at 14 dpi. Numbers above bars (plants scored); error bars, SD of three replicates. Mann–Whitney test (ns, not significant).

## 3. DISCUSSION

In this study, we identified Glu1 as a putative GH30 β-1,6-endoglucanase of the basidiomycete pathogen *U. maydis* associated with plant pathogenesis. Structural analysis showed that the substrate-binding pocket of Glu1 closely resembles that of BT3312, a biochemically characterised GH30_3 β-1,6-glucanase, but differs from that of GH30 xylanases, and that Glu1 structure is also conserved across Ustilaginales. Phylogenetic analysis also places Glu1 in GH30_3. Consistent with a catalytic role, variants of the catalytic residues largely rescued the saprophytic phenotypes of Δ*glu1* but failed to restore virulence.

### 3.1. Glu1 catalytic activity might be specifically required for virulence in Ustilaginales

Deletion of *glu1* reduces virulence, causing lower colonization of plant tissues and impaired tumour formation **(Figures 1c-d, 3f and S2b)**. Under saprophytic conditions, the Δ*glu1* mutant shows reduced septation and increased cell size **(Figures 2a and 2b)**, phenotypes fully restored by complementation with the native *glu1* allele, confirming their dependence on Glu1. Glu1 localises to the plasma membrane in close contact with the cell wall **(Figure 4a),** similar to β-1,3-glucanosyltransferases such as GAS proteins in *S. cerevisiae*, Gel1 in *A. fumigatus*, and PHR proteins in *C. albicans* (Fonzi, 1999; Mouyna et al., 2000; Mouyna et al., 2005; Ragni et al., 2007). Interestingly, complementation with Glu1 catalytic-site variants (E264A and E358A) largely restores saprophytic phenotypes but not virulence **(Figure 2 and Figure 6e)**. These findings can be explained in two, non-exclusive ways. As a first possibility, Glu1 may have a dual function: a predominantly non-enzymatic role during saprophytic growth and a catalytic function required for virulence, potentially by remodelling β-1,6-glucan or releasing β-1,6-linked oligosaccharides that could act as elicitors modulating host responses. Separation of catalytic and non-catalytic functions has been described in other organisms. In *F. oxysporum*, FoEG1 triggers plant cell death independently of its hydrolase activity, whereas its catalytic activity is required for full virulence (Zhang et al., 2021). Similarly, in the bacterium *Myxococcus xanthus*, catalytic-site variants of the cell wall enzymes PBP1a2 and DacB still retain non-catalytic functions (Zhang et al., 2023). As a second possibility, the E264A and E358A substitutions may not completely abolish catalytic activity. Since *glu1* expression peaks during colonization at 3–5 dpi (Figure 3) suggesting a much more relevant role during this growth phase, residual activity could suffice for saprophytic growth but not for infection. Distinguishing between these possibilities will require measuring the activity of the catalytic mutants.

### 3.2. Glu1 and Umag_04357 might have functional divergence in Ustilaginales fungi

Phylogenetic analysis revealed that *Ustilago maydis* encodes two GH30 family enzymes, Glu1 and Umag_04357 **(Figures 7a and 7b)**. Both proteins are structurally conserved, share the same catalytic residues **(Figures 7b and 7c)**, and are preserved across Ustilaginales **(Figures S7 and S8)**. Despite this similarity, deletion of Umag_04357 did not affect tumour formation **(Figure 7e)**. Comparable cases have been reported for other fungi: in *Aspergillus fumigatus*, the Δ*gel2* mutant shows reduced virulence, but the double mutant Δ*gel1*Δ*gel2* does not display additive defects (Mouyna et al., 2005). Similarly, in *U. maydis*, Row protein family members (Row1–Row6) share high structural conservation and are generally required for infection, except Row3 (Pejenaute-Ochoa et al., 2024).

These differences could be explained by their expression profiles. Umag_04357 is induced during infection (peak at 4 dpi), but at much lower levels than Glu1 **(Figure 7d)**. Similar patterns are observed in *Magnaporthe grisea* cutinases, where Cut2 is strongly expressed during appressorium-mediated penetration and its deletion results in severely reduced pathogenicity, whereas the other family members show lower or temporally distinct expression patterns (Skamnioti and Gurr, 2007).This indicates that possession of a catalytically active domain alone does not confer a role in infection, and that expression timing and level, rather than function, determine functional relevance during pathogenesis, a pattern mirrored by Glu1 and Umag_04357.

This divergence between Glu1 and Umag_04357 expression levels in *U. maydis* is also maintained in *Sporisorium reilianum* orthologues, where the Glu1 orthologue sr13131 is expressed nearly 1000-fold higher than the Umag_04357 orthologue sr15243 during colonization (2–4 dpi) (Zuo et al., 2021) **(Figure S9)**. Such conservation across species suggests that these differences reflect broader functional specialisation rather than host- or tissue-specific roles. Phylogenetic analyses indicate that Glu1 displays slightly longer branch lengths compared to Umag_04357 **(Figure S8)**, which could suggest a higher degree of evolutionary divergence. Although host- or tissue-specific roles cannot be discarded, these observations raise the possibility that Glu1 has acquired a more specialised role in host–pathogen interactions, whereas Umag_04357 may have retained a more basal or auxiliary function.

### 3.3. The β8–α8 insertion of Glu1 could be a lineage-specific feature of Ustilaginales

Glu1 has a predicted topology comprising a GH30 catalytic domain (residues 100–358, extended to 540 by structural alignment) and a C-terminal extension (residues 541–664). The GH30 domain contains a long insertion between β8 and α8 (residues 395–527) with a predicted intrinsically disordered region (IDR; residues 433–508) and a predicted proteolytic cleavage site (ProP) at residue 468. A second IDR is located toward the C-terminal end (residues 568–634) **(Figure 1, S1 and Table S1)**.

IDRs of the C-terminal are enriched in polar residues, which may facilitate flexible interactions with the membrane or cell wall (Macainsh et al., 2025) and act as membrane anchors in other fungal proteins such as the protein kinase Cwr1 in *Candida albicans* (Naseem et al., 2024). However, the polar content of the C-terminal IDR is not conserved across Glu1 orthologues (7–40%), and in Glu1 it is acidic (20% D+E) and lacks basic residues (3.3% K+R). It may therefore act as a flexible tail or spacer for the adjacent β-rich region, rather than as a membrane anchor.

In contrast, the IDR within the β8–α8 insertion (residues 433–508) has a similar composition in all Ustilaginales orthologues: it is enriched in charged residues (24–32%), with a modest polar content (22–28%). The insertion is much longer in Glu1 and its orthologues than in the other proteins compared (133 residues vs 37 in BT3312, 38 in MoGlu16 and 41 in Umag_04357). Since it begins close to the active site (∼10 Å from the catalytic glutamate), this flexible, charged region could facilitate substrate accessibility. Lineage-restricted features have also been reported for the smut effector Erc1, which is restricted to Ustilaginomycotina and binds plant cell wall polysaccharides (Ökmen et al., 2022). We hypothesize that this insertion represents an adaptation of this group of fungi to the modification of a more complex plant cell wall during host colonization.

The insertion also contains the predicted ProP cleavage site, which is conserved only in some Ustilaginales **(Figure S1, Table S1)**. Since this site lies between β8 and α8, cleavage there would separate α8 from the rest of the barrel and is therefore unlikely to generate a functional form. Together with the weak processing observed by western blot **(Figure 4b)**, this suggests that the ∼50 kDa band most likely reflects basal or non-specific proteolysis. Collectively, these findings link Glu1, a GH30 protein with structural similarity to GH30_3 β-(1,6)-glucanases, to virulence in a Basidiomycota plant pathogen, extending our understanding of cell wall-remodelling enzymes beyond the more extensively studied Ascomycota. The specific requirement of Glu1’s catalytic residues for infection, together with the fact that β-1,6-glucans are absent in plants and animals, positions Glu1 as an attractive and selective target for controlling smut diseases in cereal crops. Biochemical characterisation of Glu1 activity, identification of the products released from β-1,6-glucan, and structure-guided inhibitor design will be needed to fully exploit this potential and to clarify the mechanistic basis of Glu1’s role in morphogenesis and pathogenesis.

## 4. EXPERIMENTAL PROCEDURES

### 4.1. Plasmids and strain constructions

*U. maydis* strains, plasmids and primers used in this study are listed in **Tables S8 and S9**. Deletion mutants of *glu1* were generated by homologous recombination (Brachmann et al., 2004; Kämper, 2004) and of Umag_04357 by NEBuilder HiFi DNA Assembly. All deletions were verified by Southern blot and PCR. Complementation and GFP-tagging constructs were cloned into p123 (Aichinger et al., 2003) and integrated into the *ip* locus, as detailed in **Supplementary Methods (SM)**.

### 4.2. Conjugation tube, mating and filamentation assay

To assess mating and filamentation, cells were first cultured in YEPSL medium to exponential phase (OD₆₀₀ = 0.6–0.8). For solopathogenic strains (SG200), cells were adjusted to a final OD₆₀₀ of 0.8 in sterile water, and 5 μL were spotted onto Potato Dextrose (PD) charcoal plates. Plates were incubated at 25 °C for 24–48 hours. For strains of compatible mating types (FB1 and FB2), equal volumes were mixed at a 1:1 ratio prior to spotting. Successful mating or filamentation was indicated by the appearance of white, fuzzy filaments.

To assess conjugation tube formation, *U. maydis* FB1 Δ*glu1* mutant strains were cultured in liquid CMD medium to exponential phase, diluted to an OD₆₀₀ of 0.5 in 1 mL of fresh CMD and incubated at room temperature for 5 hours on a rotating wheel. Conjugation tubes were induced with 1 µL of synthetic *a2* pheromone (2.5 mg/mL).

### 4.3. Growth and cell morphology analysis

Cells were cultured in YEPSL medium to exponential phase, diluted to OD₆₀₀ = 0.05 in fresh YEPSL, and grown again to stationary phase. Growth kinetics were recorded every 15 minutes in 96-well plates under continuous shaking using a Spark 10M (Tecan) microplate reader. Experiments were performed with three independent biological replicates. For morphological and cell length analysis, cells grown to exponential phase were stained with Calcofluor White (CFW) staining solution (4 µg/mL) to visualise cell walls. Samples were examined using a DeltaVision fluorescence microscope, and cell length was quantified using ImageJ software.

### 1.1. Plant infection assays

For pathogenicity assays, *U. maydis* strains were cultured in YEPSL medium to exponential phase (OD₆₀₀ = 0.6–0.8), then adjusted to strain-specific ODs: OD₆₀₀ = 3 for CL13, OD₆₀₀ = 1 for SG200, and OD₆₀₀ = 0.5 for FB1 × FB2 crosses. Cultures were washed twice with sterile distilled water and injected into 7-day-old *Zea mays* (Early Golden Bantam) seedlings. Disease symptoms and tumour formation were scored 14 days post-inoculation. Each condition was tested in at least three independent biological replicates.

### 4.4. Fungal Biomass Analysis

For fungal biomass quantification, 2 cm long maize leaves collected at 3 days post-infection (dpi) were cut 1 cm below the inoculation site and processed as previously described (Marín-Menguiano et al., 2019). Each reaction was performed using 60 ng of total DNA as template. Fungal biomass was estimated by amplifying the constitutively expressed *ppi1* gene and normalized to the plant *gapdh* gene. Primers used are listed in **Table S8**.

### 4.5. RNA extraction and reverse transcription-quantitative PCR

For Glu1 expression analysis, total RNA was extracted from FB1 axenic cultures and from maize tissue infected with FB1 and FB2 at 1, 3, 5, and 9 days post-infection (dpi). Samples were obtained and treated as detailed in **SM**.

### 4.6. Protein and blotting assays

Total protein was obtained as described (Ochoa et al., 2026). 60 μg of each protein fraction was separated by SDS-PAGE and detected by western blot analysis as explained (Marín-Menguiano et al., 2019). Colony secretion assay was performed as previously described (Moreno-Sánchez et al., 2021).

### 4.7. Samples preparation for microscopy analysis

For appressoria formation, infected leaves were stained with CFW (0.1 mg/ml) and observed 18 h after plant inoculation. Glu1::GFP cells were immobilised on 1.8% agarose pads in PBS 1X. All samples were observed using Delta Vision. For hyphae proliferation, infected leaves from 5 dpi were stained as explained (Pejenaute-Ochoa et al., 2024) and visualised by confocal microscopy (Leica SP5 MP-AOBS). Quantification settings are detailed in **SM**.

### 4.8. Bioinformatic Analyses

Orthologues of Glu1 and Umag_04357 were retrieved by BLASTp and aligned with MAFFT L-INS-i (Katoh et al., 2018) and Jalview. Maximum-likelihood trees (JTT model, 1,000 bootstraps) were displayed in iTOL v6, and structural homologs of Glu1 were identified with AlphaFold (Varadi et al., 2024). All crystallographic structures and AlphaFold models used, as well as the predictive tools, are detailed in **SM**.

Molecular docking was carried out using the SwissDock (AutoDock Vina) web server (Bugnon et al., 2024), which employs the EADock DSS algorithm for automated ligand–protein docking. Docking parameters are detailed in **SM.**

Details Pymol alignments are given in **Tables S4-7**.

## CRediT

MDP-O, IMS, RRB and JII conceived the study. MDP-O and IMS performed the experiments and analysed the data. MDP-O wrote the original manuscript with review and editing of all the co-authors. Funding acquisition: JII and RRB.

## Acknowledgements

We would like to thank the Genetics Department of CABD for their useful discussions, Víctor Carranco, Patricia López, Clara Salmerón and Sandra Romero for their technical assistance. We thank Alfonso Fernández-Álvarez (IBFG, Salamanca, Spain), and Manuel Velasco-Gomariz, of the HIRI, Würzburg, Germany, for their careful reading of the manuscript and insightful comments. The authors used Claude (Anthropic) to assist with drafting. These tools were not used to generate or interpret scientific data or conclusions.

This research was supported by MCIN/AEI/10.13039/501100011033/ and by ‘ERDF A way of making Europe’, grant no. BIO2016-80180-P and MCIN/AEI/10.13039/501100011033/ grant no. PID2019-110477GB-I00 to JII and AEI/10.13039/501100011033/ FEDER, UE, grant no. PID2022-142783NB-I00 to JII and RRB. I.M.-S. was awarded by grant BES-2014-069149 MINECO AEI/FEDER.

## Data Availability Statement

The data that support the findings of this study are available from the corresponding author upon reasonable request.

## Supporting Information legends

**FIGURE S1.** Domain architecture of Glu1 is broadly conserved across Ustilaginales.

**FIGURE S2. (a)** Deletion of *glu1* does not impair growth under axenic conditions. **(b)** Δ*glu1* shows reduced colonization at 3 dpi.

**FIGURE S3.** Glu1 expressed under its native promoter localises to the cell periphery.

**FIGURE S4.** AlphaFold confidence of the Glu1 and Umag_04357 models.

**FIGURE S5.** Multiple sequence alignment of Glu1 orthologues and BT3312.

**FIGURE S6.** AlphaFold confidence of Glu1 orthologues.

**FIGURE S7.** Multiple sequence alignment of Glu1 and Umag_04357 orthologues.

**FIGURE S8.** Phylogenetic analysis of Glu1 and Umag_04357 orthologues reveals two conserved clades.

**FIGURE S9.** Expression of the *Sporisorium reilianum* orthologues of Glu1 and Umag_04357 during plant infection.

**TABLE S1.** Predicted features of Glu1 their orthologues across Ustilaginales and Umag_04357.

**TABLE S2.** Quantification, comparison and statistical analysis of cell morphology from Figure 2 and infection phenotypes from Figure 6e.

**TABLE S3.** Data obtained from the docking of β-gentiobiose in the Glu1 pocket

**TABLE S4.** Data obtained from structural alignment of Glu1 and its Ustilaginales orthologues with BT3312.

**TABLE S5.** Molecular docking of Glu1 against alternative substrates.

**TABLE S6.** Data obtained from the comparison of the active-site pocket of Glu1, BT3312, MoGlu16 and Xyn30B

**TABLE S7.** Equivalence of β-strands and α-helices of the (β/α)₈ barrel in Glu1 and Umag_04357.

**TABLE S8.** *U. maydis* strains used in this study.

**TABLE S9.** Plasmids and oligonucleotides used in this study.

