## Supplementary Information for "Glu1, a conserved GH30_3 β-(1,6)-glucanase requires its catalytic residues for full virulence in *Ustilago maydis*"

**The PDF file includes:**

Supplementary Materials and Methods
Supplementary Figures Legend
References

### 1. Supplementary Methods

##### Plasmids and strain constructions

*Escherichia coli* DH5α, pJET1.2/blunt (Thermo Scientific, Carlsbad, CA, United States) and pBluescript II SK (+) (Stratagene, San Diego, CA, United States) were used for cloning purposes. Deletion mutants for *glu1* (Umag_05223) were generated by homologous recombination, as described previously (Brachmann et al., 2004; Kämper, 2004). Deletion for Umag_04357 was performed using the NEBuilder HiFi DNA Assembly (New England Biolabs, Ipswich, MA United States) system. All deletions were checked by Southern blot analysis and PCR. For complementation and GFP tagging, the plasmid p123 was used (Aichinger et al., 2003). To complement the CL13Δ*glu1* strain, the construct p123:P*glu1*:*glu1* was generated via NEBuilder HiFi DNA Assembly. The *glu1* promoter and ORF were PCR-amplified from genomic DNA and cloned into p123 digested with *NotI* and *PvuII* restriction enzymes. For the catalytically inactive variant, p123:P*glu1*:*glu1(E264A_E358A)*, site-directed mutagenesis primers introducing point mutations were used with the same cloning strategy. For GFP-tagging, p123:P*glu1*:*glu1*::*gfp* was generated by amplifying the *glu1* promoter and ORF as a single fragment and cloning it into the p123 plasmid digested with *PvuII* and *NcoI* restriction enzymes. To generate p123:P*otef*:*glu1*::*gfp*, the *glu1* ORF was inserted into p123 digested with *XmaI* and *NotI* under the constitutive Potef promoter. All plasmids were linearized with *SspI* and integrated into the *ip locus* of CL13Δ*glu1* for complementation or CL13 for GFP labeling.

##### RNA extraction and reverse transcription-quantitative PCR

Samples were ground in liquid nitrogen and RNA was isolated using TRIzol reagent (Thermo Fisher Scientific) in combination with the Direct-zol RNA Miniprep Plus Kit (Zymo Research), following the manufacturers' protocols. cDNA synthesis was performed using the RevertAid H Minus First Strand cDNA Synthesis Kit (Thermo Fisher Scientific). Quantitative real-time PCR (qRT-PCR) was carried out using a CFX Connect Real-Time PCR System (Bio-Rad) and SYBR Premix Ex Taq II (Tli RNase H Plus) (Takara) to measure *glu1* transcript levels with *glu1*-specific primers. *ppi1* gene primers were used as an internal control for normalization.

#### Sequence, Structural and Phylogenetic Analysis

Phylogenetic analyses were conducted to study the evolutionary conservation of Glu1 and Umag_04357 across Ustilaginales and to place Glu1 within the GH30 family. Protein sequences of orthologs were retrieved using BLASTp. Multiple sequence alignments were generated using the L-INS-i algorithm implemented in MAFFT and visualised with Jalview to assess conservation of key residues. Phylogenetic trees were constructed using the Maximum Likelihood method based on the JTT substitution model, with initial trees obtained via the Neighbor-Joining method and 1,000 bootstrap replicates to assess branch support. Tree visualization and annotation were performed using the Interactive Tree of Life (iTOL v6) platform.

Characterized members of subfamilies GH30_3, GH30_5, GH30_7 and GH30_11 from fungi and bacteria were retrieved from the CAZy database.

| Subfamily | Protein | Organism | Accession number |
|---|---|---|---|
| GH30_3 | Glu1 (UMAG_05223) | *Mycosarcoma maydis (Ustilago maydis)* | A0A0D1E2V9 |
| GH30_3 | BT3312 | *Bacteroides thetaiotaomicron* | Q8A2J3 |
| GH30_3 | Neg1 | *Neurospora crassa* | Q7M4T0 |
| GH30_3 | LePus30A | *Lentinula edodes* | F8WPI4 |
| GH30_3 | GH30A | *Coprinopsis cinerea* | A0A7D5FSA5 |
| **GH30_5** | Bl1,6Gal | *Bifidobacterium longum* | A0A401ETL2 |
| **GH30_5** | Endo-β-1,6-galactanase | *Streptomyces avermitilis* | Q82CY3 |
| **GH30_5** | Endo-β-1,6-galactanase (NCU09702) | *Neurospora crassa* | Q7S2S4 |
| **GH30_5** | 6GAL | *Trichoderma viride* | Q76FP5 |
| GH30_7 | Xyn30B | *Talaromyces cellulolyticus* | A0A510NXC4 |
| GH30_7 | Xyn30C | *Talaromyces pinophilus* | A0A6N4SL16 |
| GH30_7 | Xyn30A | *Thermothelomyces thermophilus* | G2Q1N4 |
| GH30_7 | Neg1-like (β-1,6-glucanase annotation) | *Sugiyamaella lignohabitans* | A0A161HKM6 |
| **GH30_11** | BgalA | *Fomitopsis palustris* | A0A5B8GWC9 |
| **GH30_11** | Glycoside hydrolase | *Stereum hirsutum* | EIM85030.1 |
| **GH30_11** | Glycoside hydrolase | *Trametes versicolor* | EIW53353.1 |
| Outgroup | β-1,3-Endoglucanase EglC (GH17) | *Aspergillus fumigatus* | EAL86311.1 |

**Protein structures and structural alignment**

Predicted 3D structures of Glu1 (UniProt A0A0D1E2V9), Umag_04357 (UniProt A0A0D1CK77), MoGlu16 (UniProt G4MNS0) from *Magnaporthe oryzae* and Glu1 orthologs: *P. hubeiensis* (UniProt R9NWT8), *S. reilianum* (UniProt E6ZYW6), *U. bromivora* (UniProt A0A1K0G2P4) and *M. pennsylvanicum* (UniProt A0A077R274), were obtained from the AlphaFold Protein Structure Database (model version 6 for all entries). The crystal structures of BT3312 (PDB 5NGL) and Xyn30B (PDB 6IUJ), both solved by X-ray crystallography, were used for the structural comparisons.

**Predictive analysis tools**

Protein sequences were retrieved from the UniProt database. Conserved domains were identified using InterPro (Blum et al., 2021) . Signal peptides were predicted using SignalP v6.0 (Teufel et al., 2022) . N- and O-glycosylation sites were predicted with NetNGlyc 1.0 (Gupta and Brunak, 2002) and NetOGlyc 4.0 (Steentoft et al., 2013) . Effector properties and GPI-anchor signals were predicted using EffectorP-fungi 6.0 (Sperschneider and Dodds, 2022) respectively. Proteolytic cleavage sites were assessed with ProP 1.0 (Duckert et al., 2004) ). Protein topology was represented using IBS 2.0 (Xie et al., 2022) .

**Molecular docking**

The predicted 3D structure of Glu1 was retrieved from the AlphaFold Protein Structure Database (Varadi et al., 2024) in PDB format. Ligand structures were obtained in SMILES format from PubChem (CID reported in **Table S2**). Molecular docking was carried out using the SwissDock (AutoDock Vina) web server (Bugnon et al., 2024) , which employs the EADock DSS algorithm for automated ligand–protein docking. AutoDock Vina was performed with a sampling exhaustivity value of 40. The docking search space was centered on the predicted catalytic site based on structural modeling, using a box size of 20 × 20 × 20 Å and a box center at coordinates (-3, 8, -3) for Glu1 and (1, 6, 18) for BT3312. Docking poses were ranked based on FullFitness scores and estimated binding free energies (ΔG). The resulting clusters were visualised and analysed using PyMOL.

##### Statistical analysis

Statistical analyses were performed in GraphPad Prism v10. Symptoms scoring of infected plants **(Figures 1c–e, 6e and 7e)** were compared pairwise using the Mann–Whitney test. Septum number and cell size **(Figure 2)** were compared among strains using the one-way ANOVA Kruskal–Wallis test followed by Dunn's post-hoc test for multiple comparisons. Gene expression, conjugation tube formation, appressorium formation, quantification of fungal colonization from confocal microscopy images **(Figure 3)** and fungal biomass **(Figure S2b)** were analysed with Student's t-test. $P < 0.05$ was considered significant (*$P < 0.05$; **$P < 0.005$; ***$P < 0.0005$; ****$P < 0.0001$; ns, not significant). Unless otherwise stated, error bars represent the standard deviation (SD) of two/three independent biological replicates. Sample sizes (plants or cells scored), replicate numbers, pairwise comparisons and exact tests are given in the figure legends and in **Table S2**.

##### Use of artificial intelligence (AI) tools

The authors used Claude (claude-sonnet-5-5, Anthropic) to help write PyMOL scripts for the structural analyses and to assist with drafting and English language editing. All outputs were reviewed by the authors. AI tools were not used to generate or interpret experimental data.

#### 2. Supporting Information legends

**FIGURE S1. Domain architecture of Glu1 is broadly conserved across Ustilaginales.** Schematic predicted domain architecture of Glu1 and its orthologs in representative Ustilaginales and related Basidiomycota (*Sporisorium reilianum*, *S. scitamineum*, *Ustilago hordei*, *U. bromivora*, *Moesziomyces antarcticus*, *Pseudozyma hubeiensis*, *Melanopsichium pennsylvanicum* and *Testicularia cyperi*). Signal peptide (SP) in lilac, GH30 domain in pink, intrinsically disordered regions (IDR) in yellow, N- and O-glycosylation sites as blue hexagons and green circles and predicted processing sites as teal arrows. The positions of the catalytic glutamates are indicated by a star, and protein length (aa) is at the right of each ortholog.

**FIGURE S2. Deletion of *glu1* does not impair growth under axenic conditions.** a) Growth curves of CL13 and Δ*glu1* strains in liquid YEPSL, measured as optical density (O.D., 595 nm) over 20 h. Error bars, SD of three independent replicates. b) Quantification of fungal biomass *in planta*, estimated by *ppi1* expression relative to the plant *gapdh* gene in CL13 (WT), Δ*glu1* and mock-inoculated leaves at 5 dpi. Error bars, SD of three independent replicates; Student’s t-test (P = 0.1).

**FIGURE S3. Glu1 expressed under its native promoter localizes to the cell periphery.** Representative brightfield and GFP images expressing P*glu1*:*glu1*::*gfp* under the control of the endogenous *glu1* promoter, grown under saprophytic conditions (scale bar, 10 µm).

**FIGURE S4. AlphaFold confidence of the Glu1 and Umag_04357 models.** Predicted aligned error (PAE) plots and mean pLDDT of the AlphaFold models of Glu1 (81.25) and Umag_04357 (87.65).

**FIGURE S5. Multiple sequence alignment of Glu1 orthologs and BT3312.** Alignment of Glu1 orthologs across Ustilaginales and *Bacteroides thetaiotaomicron* BT3312 (top sequence), obtained with MAFFT v7 and displayed in Jalview with the Clustal colour scheme. Conservation, quality and consensus tracks are shown below the alignment.

**FIGURE S6. AlphaFold confidence of Glu1 orthologs.** PAE plots and mean pLDDT of the AlphaFold models of *P. hubeiensis* (AF-R9NWT8, 74.69), *S. reilianum* (AF-E6ZYW6-F1, 77.62), *U. bromivora* (AF-A0A1K0G2P4, 82) and *M. pennsylvanicum* (AF-A0A077R274, 83.06).

**FIGURE S7. Multiple sequence alignment of Glu1 and Umag_04357 orthologs.** Alignment underlying the comparison shown in Figure 7c, obtained with MAFFT v7 and displayed in Jalview. Glu1 orthologs (blue) and Umag_04357 orthologs (green) are indicated at the left. Conservation, quality, consensus and occupancy tracks are shown below each block.

**FIGURE S8. Phylogenetic analysis of Glu1 and Umag_04357 orthologs reveals two conserved clades.** Maximum-likelihood tree of Glu1 (blue star) and Umag_04357 (green star) orthologs across Ustilaginales, rooted with the outgroup (red star). Homologs were identified with BLASTp and aligned with MAFFT v7; the tree was generated with Archaeopteryx.js and edited in iTOL. Branch lengths are shown above each branch (scale bar, 1 substitution per site); nodes with bootstrap support >90 (1,000 replicates) are marked with a blue circle.

**FIGURE S9. Expression of the *Sporisorium reilianum* orthologs of Glu1 and Umag_04357 during plant infection.** Expression profiles based on the RNA-seq data from Zuo et al. (2021) for the *S. reilianum* orthologs of Glu1 (sr13131) and Umag_04357 (sr15243) in axenic culture (ax) and post-infection (dpi) conditions. **(a)** Expression of sr13131, the Glu1 ortholog. **(b)** Expression of sr15243, the Umag_04357 ortholog. **(c)** Comparison of both genes on the same scale across the same time course.

**TABLE S1. Predicted features of Glu1 and Umag_04357 and their orthologs across Ustilaginales.** Boundaries of the signal peptide, GH30 domain and intrinsically disordered regions, catalytic residue positions, predicted proteolytic processing sites and N- and O-glycosylation sites, corresponding to the domain architectures in Figures 1a, 7a and S1.

**TABLE S2. Quantification, comparison and statistical analysis of cell morphology and infection phenotypes.** Quantification of cell length, septation and disease symptoms in WT, Δ*glu1*, complemented and catalytic-site variant (E264A/E358A) strains (Figures 2 and 6e), with sample sizes, pairwise comparisons between strains and the statistical tests used (Mann–Whitney; $p < 0.05$ considered significant).

**TABLE S3. Data obtained from structural alignment of Glu1 and its Ustilaginales orthologs with BT3312.** RMSD, number of aligned residues, GH30 domain boundaries (annotated and structurally extended), position and length of the β8–α8 insertion, and secondary structure assignment of the eight β-strands and α-helices of the $(\beta/\alpha)_8$ barrel. The Pymol settings used are detailed in the table file.

**TABLE S4. Data obtained from the docking of β-gentiobiose in the Glu1 pocket.** Residues within 4.5 Å of the ligand, polar contacts, distances to the catalytic residues Glu264 and Glu358, and displacement and RMSD between the three poses. The docking was performed with SwissDock, and the PyMOL settings used for the analysis are detailed in the table file.

**TABLE S5. Data obtained from the comparison of the active-site pocket of Glu1, BT3312, MoGlu16 and Xyn30B.** Equivalent residues in loops 1 and 2, catalytic residues and subsites, with Cα–Cα distances and RMSD from the structural alignments. The PyMOL settings used are detailed in the table file.

**TABLE S6. Molecular docking of Glu1 against alternative substrates.** Predicted binding free energies (ΔG, kcal/mol) for β-gentiobiose, β-1,4-xylan, β-1,3-glucan and chitin-derived oligomers in the Glu1 pocket, and for β-gentiobiose in BT3312.

**TABLE S7. Equivalence of β-strands and α-helices of the $(\beta/\alpha)_8$ barrel in Glu1 and Umag_04357.** Residue ranges of each secondary structure element and length of the β8–α8 segment in both proteins.

**TABLE S8. *U. maydis* strains used in this study.**

**TABLE S9. Plasmids and oligonucleotides used in this study.** Plasmid constructs, cloning strategy used for each strain and primer sequences.

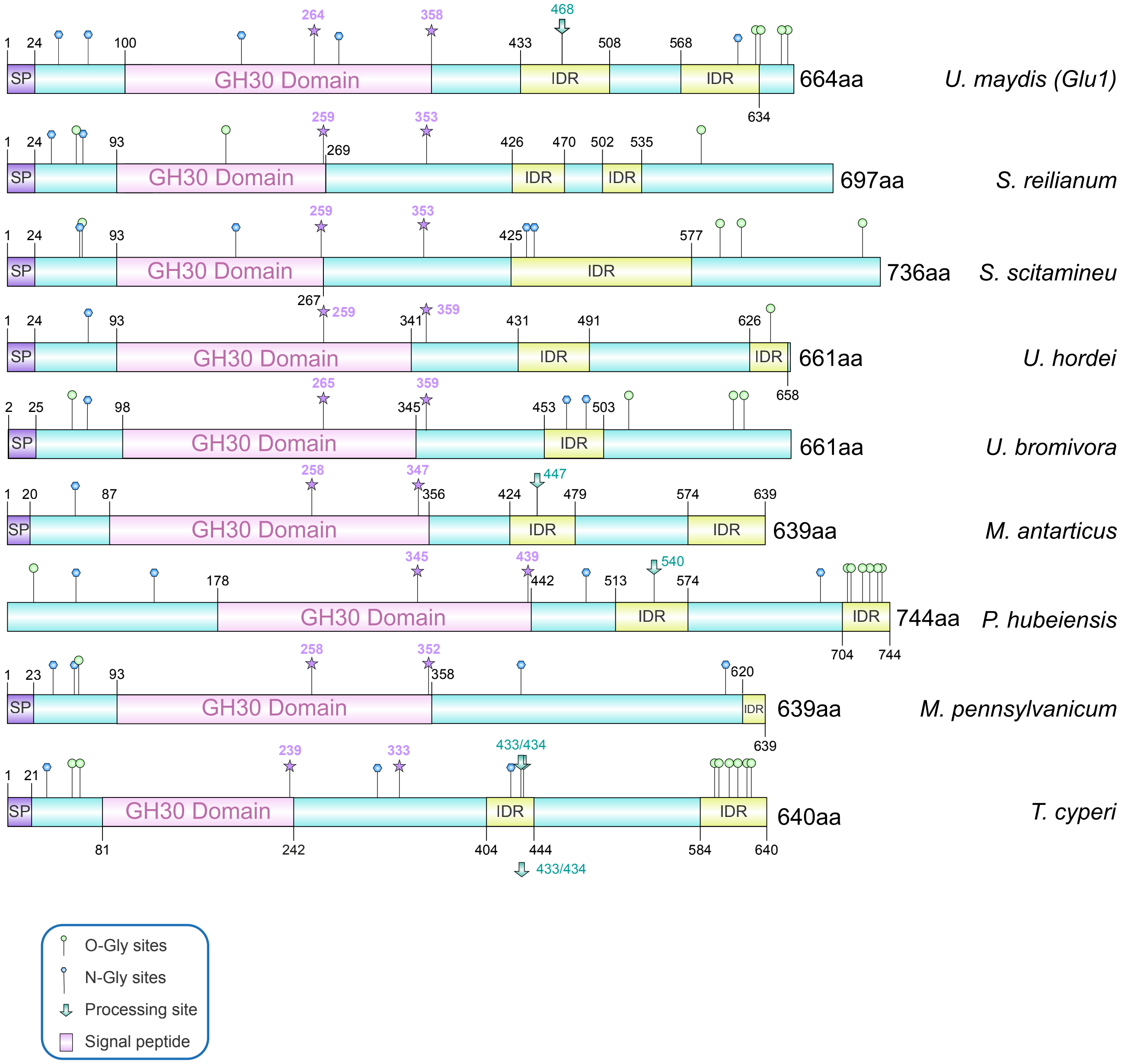

U. maydis (Glu1)
664aa
S. reilianum
697aa
S. scitamineu
736aa
U. hordei
661aa
U. bromivora
661aa
M. antarticus
639aa
P. hubeiensis
744aa
M. pennsylvanicum
639aa
T. cyperi
640aa
SP
GH30 Domain
IDR
O-Gly sites
N-Gly sites
Processing site
Signal peptide

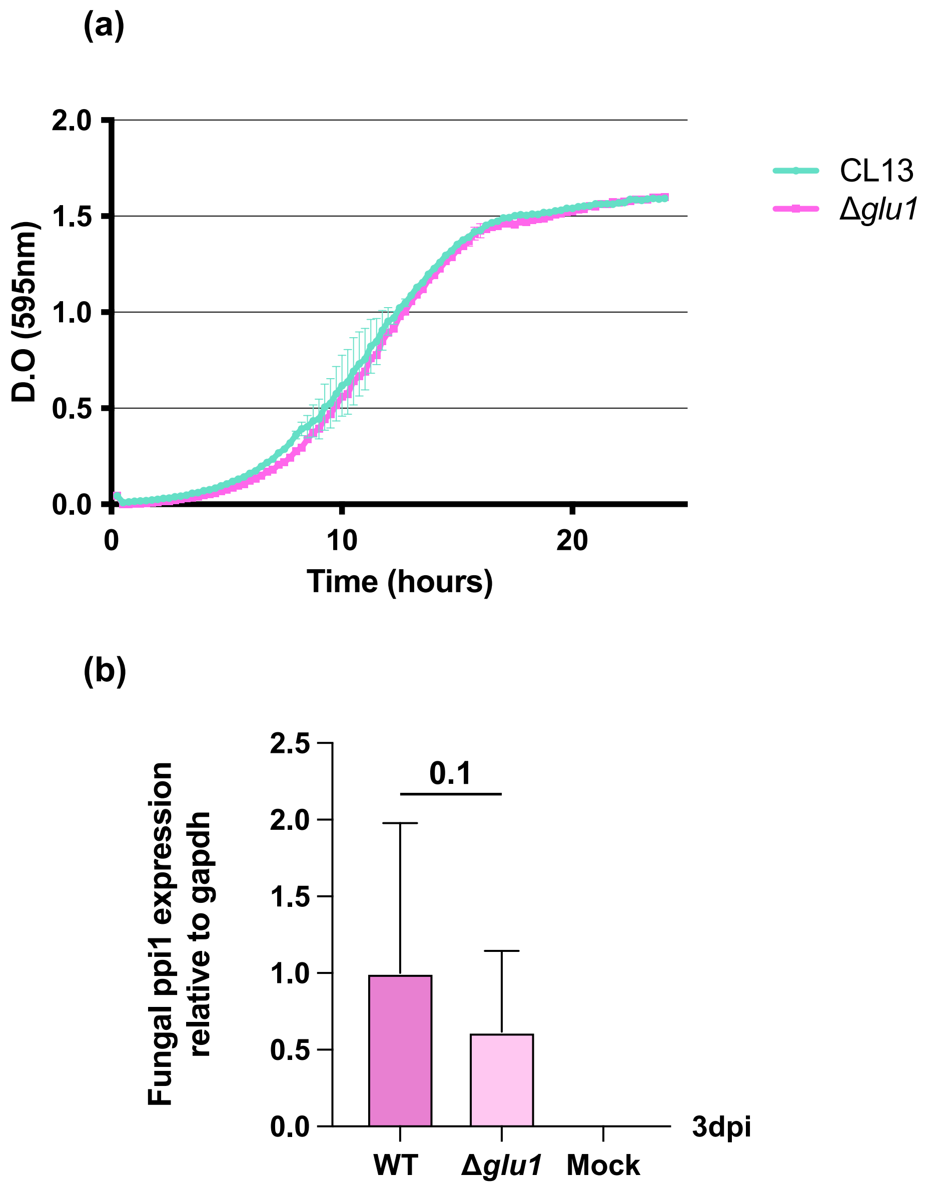

(a)
2.0
1.5
1.0
0.5
0.0
D.O (595nm)
0
10
20
Time (hours)
CL13
Δglu1
(b)
2.5
2.0
1.5
1.0
0.5
0.0
0.1
Fungal ppi1 expression
relative to gapdh
3dpi
WT
Δglu1
Mock

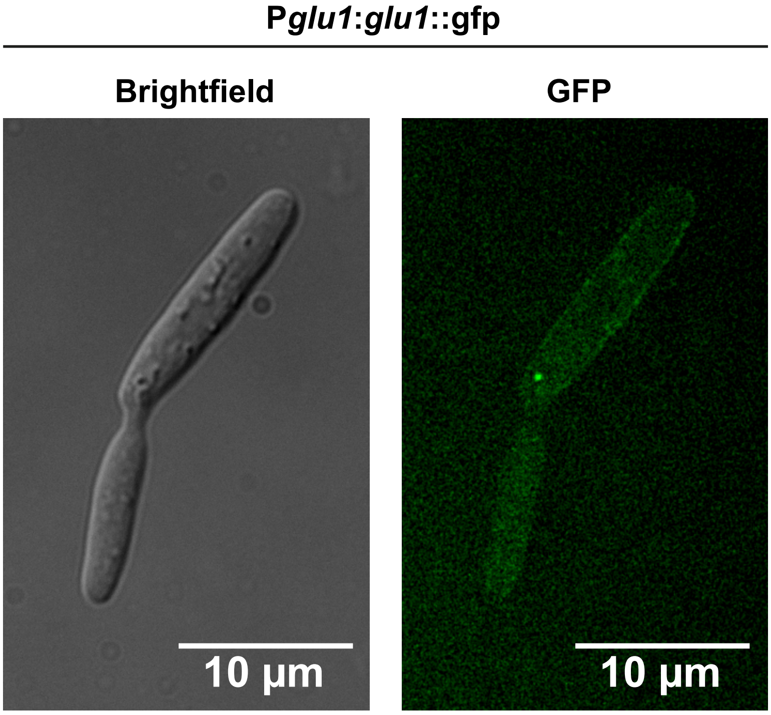
P*glu1*:*glu1*::gfp
Brightfield
GFP
10 µm
10 µm

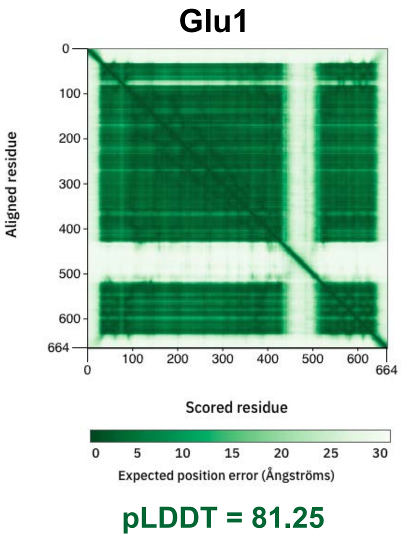
Glu1
Aligned residue
Scored residue
Expected position error (Ångströms)
pLDDT = 81.25

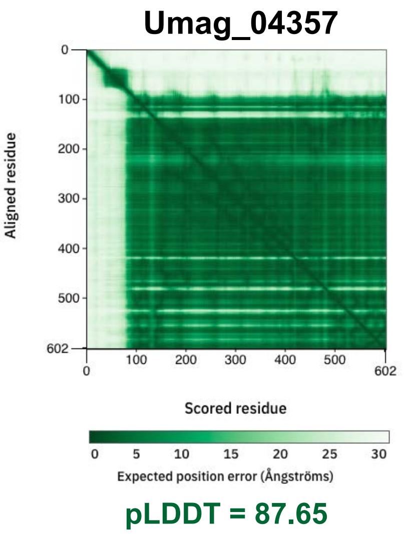
Umag_04357
Aligned residue
Scored residue
Expected position error (Ångströms)
pLDDT = 87.65

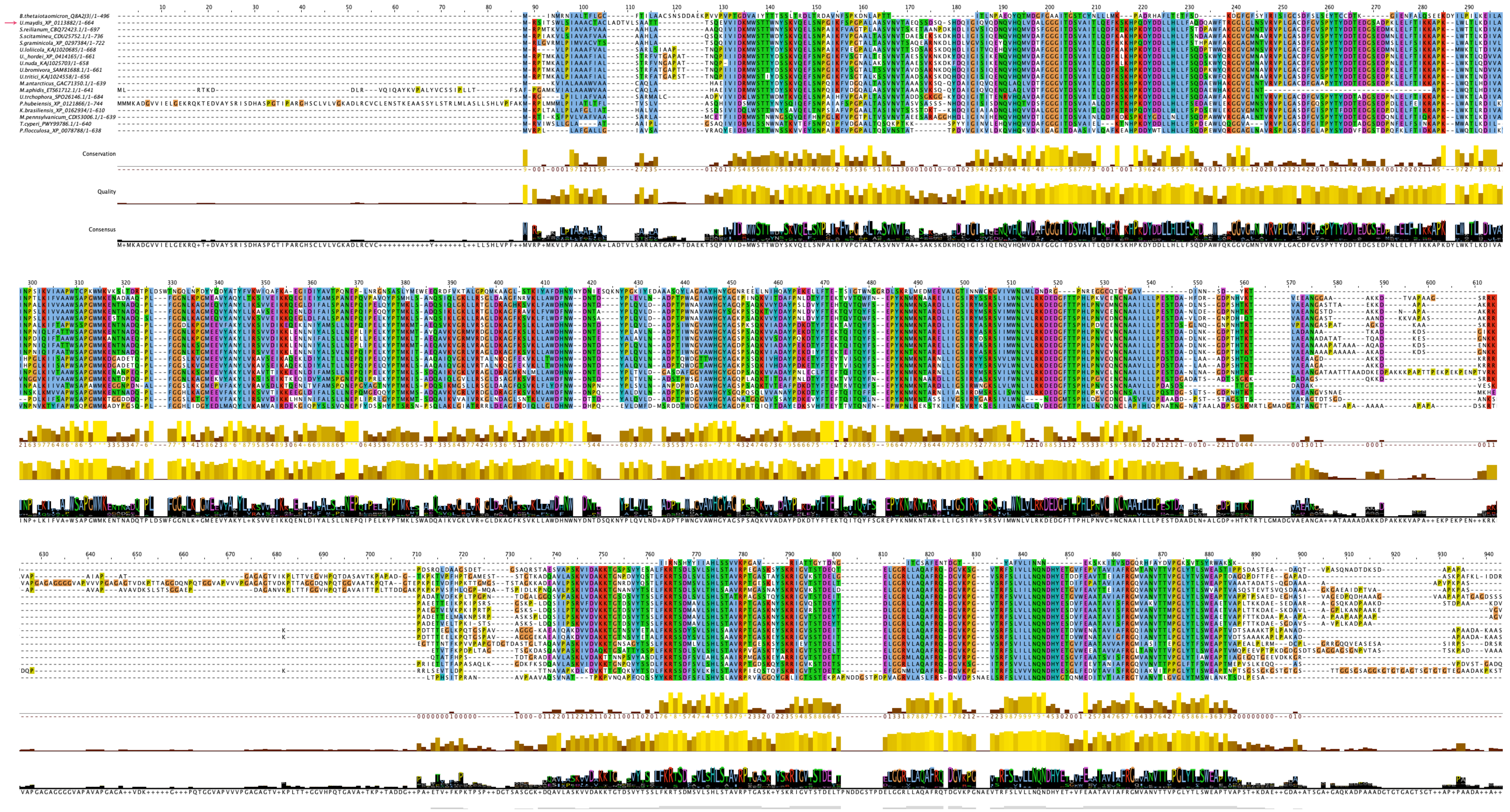
B.thetaiotaomicron_Q8A2J3|/1-496
U.maydis_XP_0113882/1-664
S.reilianum_CBQ72423.1/1-697
S.scitamineu_CDU25752.1/1-736
S.graminicola_XP_0297384/1-722
U.loliicola_KAJ1020685/1-668
U._hordei_XP_0414165/1-661
U.nuda_KAJ1025703/1-658
U.bromivora_SAM81688.1/1-661
U.tritici_KAJ1024558/1-656
M.antarcticus_GAC71350.1/1-639
M.aphidis_ETS61712.1/1-641
U.trchophora_SPO26146.1/1-684
P.hubeiensis_XP_0121866/1-744
K.brasiliensis_XP_0162934/1-610
M.pennsylvanicum_CDI53006.1/1-639
T.cyperi_PWY99786.1/1-640
P.flocculosa_XP_0078788/1-638
Conservation
Quality
Consensus

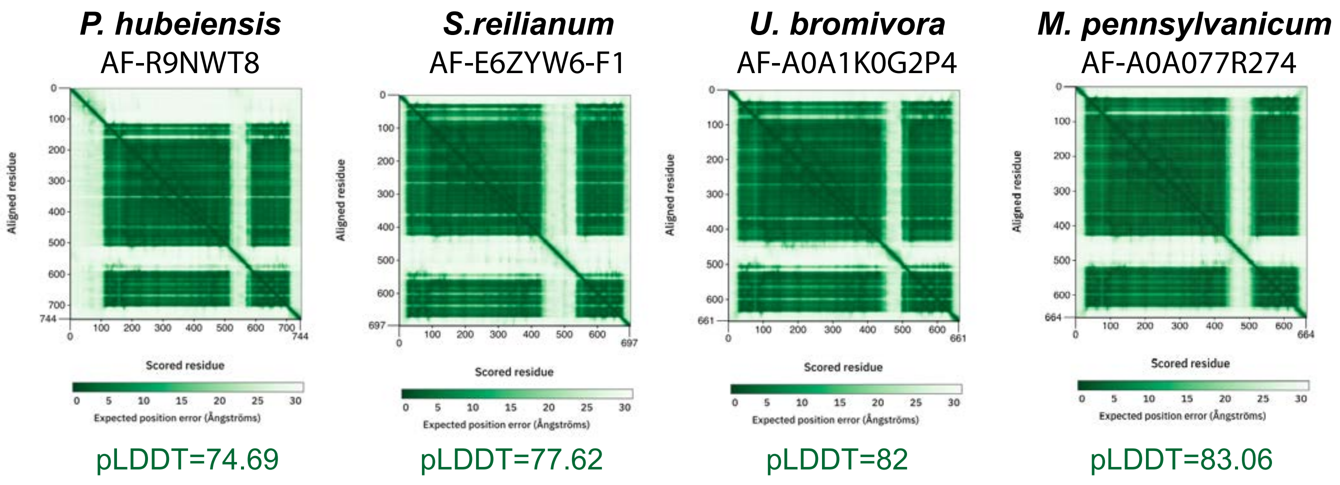

*P. hubeiensis*
AF-R9NWT8
*S.reilianum*
AF-E6ZYW6-F1
*U. bromivora*
AF-A0A1K0G2P4
*M. pennsylvanicum*
AF-A0A077R274
Aligned residue
Scored residue
Expected position error (Ångströms)
pLDDT=74.69
pLDDT=77.62
pLDDT=82
pLDDT=83.06

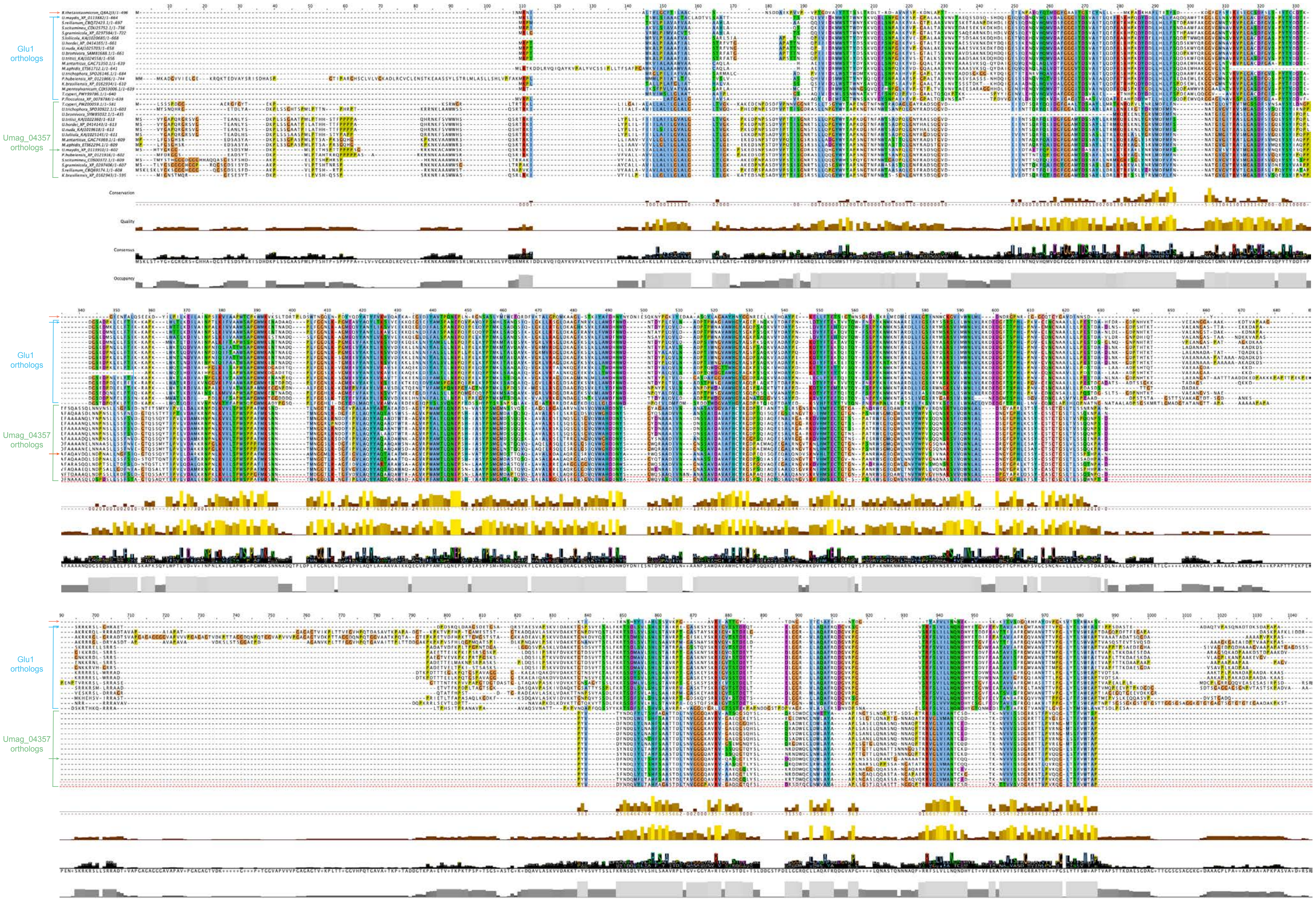
Glu1 orthologs
Umag_04357 orthologs
Conservation
Quality
Consensus
Occupancy
Glu1 orthologs
Umag_04357 orthologs
Glu1 orthologs
Umag_04357 orthologs

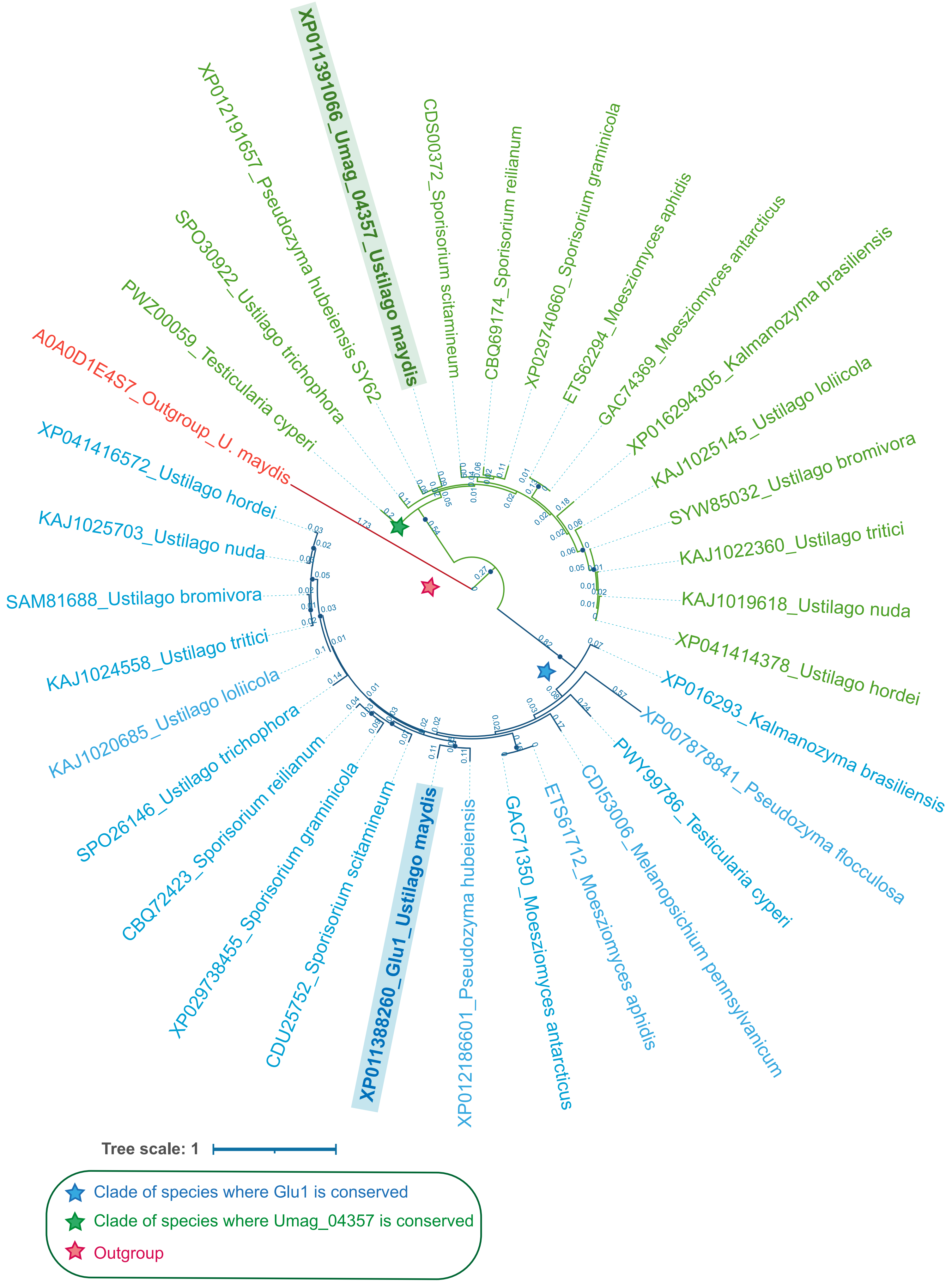

XP011391066_Umag_04357_Ustilago maydis
XP012191657_Pseudozyma hubeiensis SY62
CDS00372_Sporisorium scitamineum
CBQ69174_Sporisorium reilianum
XP029740660_Sporisorium graminicola
ETS62294_Moesziomyces aphidis
GAC74369_Moesziomyces antarcticus
XP016294305_Kalmanozyma brasiliensis
KAJ1025145_Ustilago loliicola
SYW85032_Ustilago bromivora
KAJ1022360_Ustilago tritici
KAJ1019618_Ustilago nuda
XP041414378_Ustilago hordei
SPO30922_Ustilago trichophora
PWZ00059_Testicularia cyperi
A0A0D1E4S7_Outgroup_U. maydis
XP041416572_Ustilago hordei
KAJ1025703_Ustilago nuda
SAM81688_Ustilago bromivora
KAJ1024558_Ustilago tritici
KAJ1020685_Ustilago loliicola
SPO26146_Ustilago trichophora
CBQ72423_Sporisorium reilianum
XP029738455_Sporisorium graminicola
CDU25752_Sporisorium scitamineum
XP011388260_Glu1_Ustilago maydis
XP012186601_Pseudozyma hubeiensis
GAC71350_Moesziomyces antarcticus
ETS61712_Moesziomyces aphidis
CDI53006_Melanopsichium pennsylvanicum
PWY99786_Testicularia cyperi
XP007878841_Pseudozyma flocculosa
XP016293_Kalmanozyma brasiliensis
Tree scale: 1
Clade of species where Glu1 is conserved
Clade of species where Umag_04357 is conserved
Outgroup

(a)

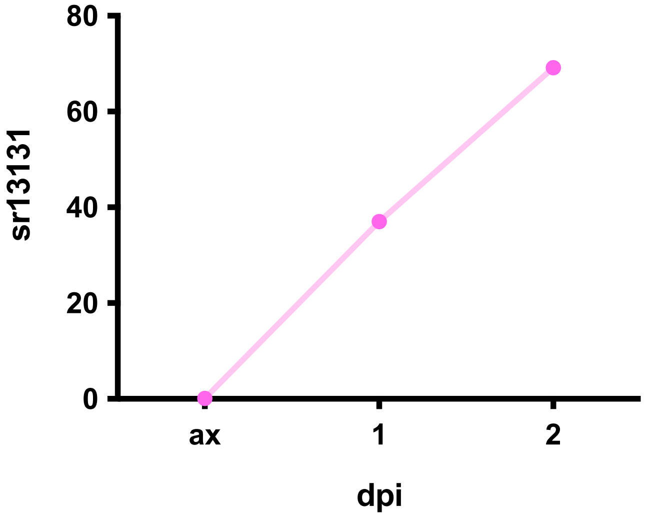

80
60
40
20
0
sr13131
ax
1
2
dpi


(b)

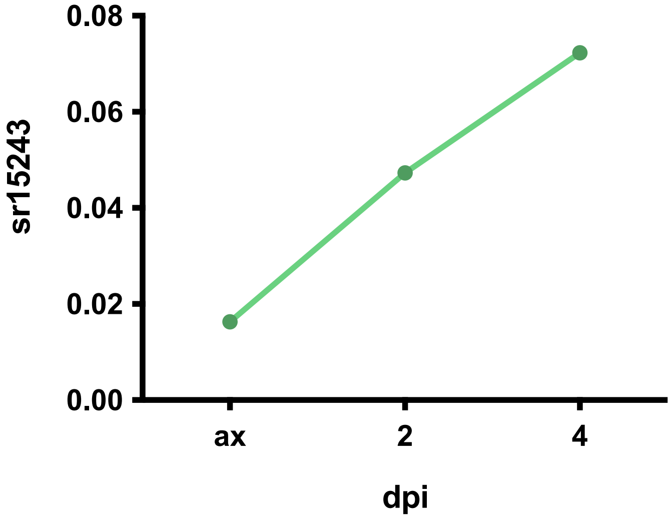

0.08
0.06
0.04
0.02
0.00
sr15243
ax
2
4
dpi


(c)

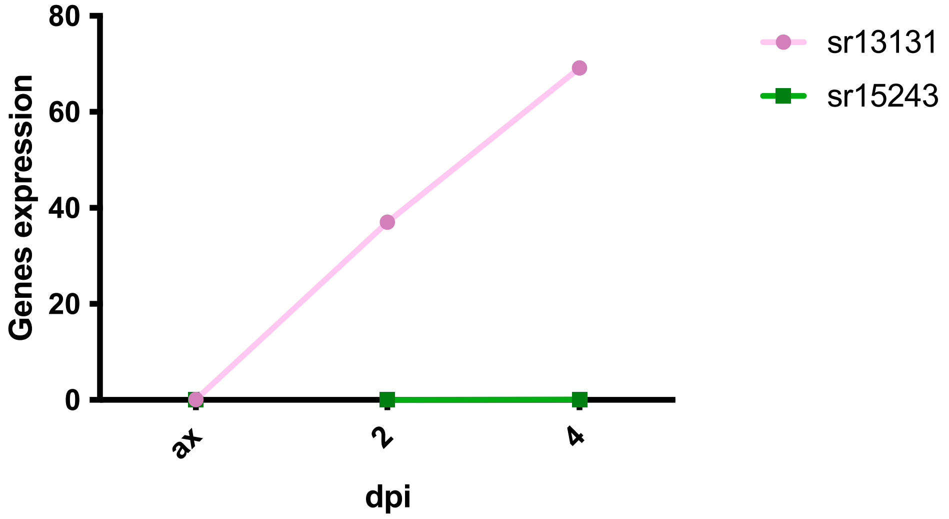

80
60
40
20
0
Genes expression
ax
2
4
dpi
sr13131
sr15243

#### Table S1: Predicted features of Glu1 and Umag_04357 and their orthologs across Ustilaginales

*Boundaries (amino acid positions) of the signal peptide, GH30 domain and intrinsically disordered regions, catalytic residue positions, predicted proteolytic processing sites (ProP) and N- and O-glycosylation sites, corresponding to the domain architectures in Figures 1a, 7a and S1.*

| Protein | Length | EffectorP (6.0-Fungi) | SignalP 6.0 | GH30 Domain Predicted | Gly Domain | DISORDER DOMAIN | Catalytic residues | ProP | GPI Anchor site | N-GLY | O-GLY (>0.7) |
|---|---|---|---|---|---|---|---|---|---|---|---|
| Umag_05223 (Glu1)_*U. maydis* | 664 | No (0.85) | 1-24aa | 100-358aa | 95-435 | 433-508<br>568-634 | 264 / 358 | 468 | No | 44, 69, 198, 280, 616 | 630, 631, 636, 640, 658 |
| CBQ72423.1_*S.reilianum* | 697 | No (0.85) | 1-24aa | 93-269 | 260, 353 | 426-470<br>502-535 | 259 / 353 | No | No | 38, 63 | 61, 581, 584 |
| CDU25752.1_*S. scitamineum* | 736 | No (0.85) | 1-24aa | 93-267 | 90-430 | 425-577 | 259 / 353 | No | No | 63, 193, 434, 443 | 61, 600, 622, 721 |
| XP_041416572.1_ *U. hordei* | 661 | No (0.88) | 1-24aa | 99-341 | 96-436 | 432-492<br>629-661 | 265 / 359 | No | No | 69 | 643 |
| SAM81688.1_*Ustilago bromivora* | 661 | No (0.81) | 1-24aa | 98-345 | 96-437 | 451-501 | 265 / 359 | No | No | 69, 471, 488 | 56, 524, 617, 622 |
| GAC71350.1_Moesziomyces antarcticus T-34 | 639 | No (0.88) | 1-20aa | 87-356 | 51-426 | 424-479<br>620-639 | 258 / 347 | 447 | No | 58 | |
| XP_012186601.1_*Pseudozyma hubeiensis SY62* | 744 | No (0.85) | No | 178-442 | 176-504 | 513-574<br>704-744 | 345 / 439 | 540 | No | 59, 125, 150, 685 | 152, 710, 719, 720, 727, 732, 734, 727 |
| CDI53006.1_*Melanopsichium pennsylvanicum 4* | 639 | No (0.94) | 1-23aa | 93-358 | 89-430 | 620-639 | 258 / 352 | No | No | 37, 62, 433, 601 | 60 |
| PWY99786.1_*Testicularia cyperi* | 640 | No (0.79) | 1-21aa | 81-242 | 79-409 | 404-444<br>584-640 | 239 / 333 | 433; 434 | No | 34, 313, 426 | 57, 61, 594, 598, 599, 604, 605, 608, 623 |
| PWN53486.1_*Violaceomyces* palustris | 577 | No (0.77) | No | 76-246 | 71-404 | No | 239 | 433; 434 | No | 51, 314, 546 | 42, 54 |
| Umag_04357_*U. maydis* | 602 | No (0.92) | No | 150-465 | 148-499 | 1-39 | 317 / 413 | 74 | No | 85, 114, 119, 186, 375, 544 | 27, 122 |

### Table S2

*Quantification, comparison and statistical analysis of cell morphology from Figure 2 and infection phenotypes from Figure 6e.*

#### Mann-Whitney statistical analysis (Fig 2a)

| | Δ*glu1* | P*glu1:glu1* | P*glu1:glu1* (E264A/E358A) |
|---|---|---|---|
| WT | 0.0003 (***) | ns | ns |
| Δ*glu1* | | 0.074 (*) | 0.006 (**) |
| P*glu1:glu1* | | | ns |

#### Kruskal–Wallis one-way ANOVA (Fig 2b)

| | Δglu1 | Pglu1:glu1 | P*glu1:glu1* (E264A/E358A) |
|---|---|---|---|
| WT | <0.0001 (****) | ns | ns |
| Δ*glu1* | | <0.0001 (****) | <0.0001 (****) |
| P*glu1:glu1* | | | ns |

#### Mann-Whitney statistical analysis (Fig 6f)

| | Δ*glu1* | P*glu1:glu1* | P*glu1:glu1* (E264A/E358A) |
|---|---|---|---|
| WT | 0.016 (**) | ns | <0.0001 (****) |
| Δ*glu1* | | 0.0034 (**) | ns |
| P*glu1:glu1* | | | 0.0039 (**) |

### Table S3: Glu1 orthologs: GH30 domain boundaries and secondary structure of the (β/α)$_8$ barrel compared with BT3312

**Structures used:**

***Ustilago maydis* (UM):** AF- A0A0D1E2V9-F1-model_v6 AlphaFold model

***Sporisorium reilianum* (SR):** AF- E6ZYW6-F1-model_v6 AlphaFold model

***Ustilago bromivora* (UB):** AF-A0A1K0G2P4-F1-model_v6 AlphaFold model

***Melanopsichium pennsylvanicum* (MP):** AF-A0A077R274-F1-model_v6 AlphaFold model

***Pseudozyma hubeiensis* (PH):** AF-R9NWT8-F1-model_v6 AlphaFold model

**BT3312:** crystal structure 5NGL (chain A)

**Table 1. GH30 domain predicted for each protein, core defined in this study and structural alignment with BT3312 and Glu1**

| Protein | Length (aa) | Predicted GH30 domain | Core defined in this study | Residues in core | RMSD vs BT3312 (Å) | RMSD vs Glu1 (Å) |
|---|---|---|---|---|---|---|
| **BT3312 (5NGL)** | 496 | 82-427 (Temple et al 2017) | 82–427 | 346 | – | 1.07 (242) |
| **Glu1 (UM)** | 664 | 100–358 | 100–540 | 441 | 1.07 (242) | – |
| **SR** | 697 | 93–269 | 93–576 | 484 | 0.96 (203) | 0.33 (315) |
| **UB** | 661 | 98–345 | 98–538 | 441 | 1.04 (236) | 0.33 (316) |
| **MP** | 639 | 93–358 | 93–523 | 431 | 0.99 (206) | 0.37 (329) |
| **PH** | 744 | 178–442 | 178–610 | 433 | 1.00 (228) | 0.23 (300) |

***Table 1:*** *Predicted GH30 domains of Glu1 orthologs in Ustilaginales. The core spans from the start of the predicted domain to the end of the eighth α-helix (α8) identified by structural alignment with BT3312. RMSD values (Å) were calculated on Cα atoms of the cores with* ***PyMOL align*** *(up to 10 cycles of outlier rejection, cutoff 2.0 Å); aligned Cα atoms are in parentheses.*

**Table 2. Equivalent β-strands and α-helices of the barrel and catalytic glutamates**

| Element | BT3312 | Glu1 | SR | UB | MP | PH |
|---|---|---|---|---|---|---|
| **β1** | 83–87 | 99–103 | 94–98 | 100–105 | 93–97 | 180–184 |
| **β2** | 122–126 | 147–151 | 142–146 | 148–152 | 141–145 | 228–232 |
| **β3** | 173–177 | 202–207 | 197–201 | 203–207 | 196–200 | 283–287 |
| **β4** | 232–234 | 258–260 | 253–255 | 259–262 | 252–255 | 339–341 |
| **β5** | 276–281 | 303–308 | 298–304 | 304–309 | 297–302 | 384–389 |
| **β6** | 310–314 | 329–333 | 324–328 | 330–334 | 323–327 | 410–414 |
| **β7** | 334–337 | 353–356 | 348–351 | 354–357 | 347–350 | 434–437 |
| **β8** | 373–376 | 391–394 | 386–389 | 392–394 | 385–388 | 472–474 |
| **Catalytic acid/base (β4)** | Glu238 | Glu264 | Glu259 | Glu265 | Glu258 | Glu345 |
| **Catalytic nucleophile (β7)** | Glu339 | Glu358 | Glu353 | Glu359 | Glu352 | Glu439 |

| Element | BT3312 | Glu1 | SR | UB | MP | PH |
|---|---|---|---|---|---|---|
| **α1a** | 90–98 | 106–118 | 101–113 | 107–119 | 100–112 | 187–199 |
| **α1b** | 101–112 | 120–131 | 115–126 | 121–132 | 114–125 | 201–212 |
| **α2** | 152–168 | 187–197 | 182–192 | 186–198 | 179–191 | 266–278 |
| **α3** | 209–226 | 233–252 | 228–247 | 234–253 | 227–246 | 314–333 |
| **α4** | 252–270 | 279–296 | 274–291 | 280–297 | 273–290 | 360–377 |
| **α5** | 294–300 | 315–326 | 310–321 | 316–327 | 309–320 | 396–408 |
| **α6** | 320–329 | 339–348 | 334–343 | 340–349 | 333–342 | 420–429 |
| **α7** | 351–369 | 370–386 | 365–381 | 371–387 | 364–381 | 451–467 |
| **Segment β8→α8** | 377–413 (37) | 395–527 (133) | 390–565 (176) | 395–526 (132) | 389–512 (124) | 475–596 (122) |
| **α8** | 414–425 | 528–540 | 566–578 | 528–540 | 513–525 | 597–609 |

***Table 2. Equivalence of β-strands and α-helices of the (β/α)$_8$ barrel in BT3312 and its homologs.*** *For each BT3312 strand or helix, the segment of the same type with the closest Cα centroid in the homolog superimposed on BT3312 with cealign was selected. Residue ranges in blue correspond to elements not detected by DSS. For these, the equivalent segment was identified from the residues within 3.5 Å of the BT3312 element after superposition and confirmed by the Cα(i)–Cα(i+3) distances (≈9–10 Å in strands and ≈5–6 Å in helices).. Values in parentheses in the β8→α8 row are the number of residues between β8 and α8. In all homologs, the eighth helix lies beyond an insertion of 122–176 residues absent from BT3312.*

### Table S4: Glu1 residues in the gentiobiose-binding pocket predicted by VINA-docking

**Residues within 4.5 Å of β-gentiobiose in the three Glu1 docking poses, distances from Cα, side chain and main chain, and pLDDT.**

##### Pose 1

| Residue | Cα–ligand (Å) | Side chain–ligand (Å) | Main chain–ligand (Å) | Polar contact (≤3.2 Å) | pLDDT (Cα) |
|---|---|---|---|---|---|
| **Glu264** | 6.9 | 2.7 (OE2–O) | 6.9 (CA–O) | Side chain | 98.4 |
| **Glu358** | 7.1 | 2.8 (OE1–O) | 7.1 (CA–O) | Side chain | 98.6 |
| Tyr335 | 3.8 | 3.2 (OH–O) | 2.9 (O–O) | Side and main chain | 97.5 |
| Thr360 | 4.9 | 4.5 (CG2–O) | 3.0 (O–O) | Main chain | 95.8 |
| Trp395 | 6.2 | 3.1 (NE1–O) | 6.2 (CA–O) | Side chain | 98.0 |
| Asp156 | 7.0 | 3.5 (OD2–C) | 7.0 (CA–C) | No | 96.8 |
| Val361 | 3.6 | 4.3 (CG1–O) | 3.6 (CA–O) | No | 92.7 |
| Val362 | 4.7 | 3.9 (CG1–O) | 3.7 (N–O) | No | 89.8 |
| Tyr272 | 10.0 | 4.3 (OH–C) | 10.0 (CA–C) | No | 95.4 |
| Phe308 | 7.9 | 4.5 (CE2–O) | 7.9 (CA–O) | No | 98.7 |

##### Pose 2

| Residue | Cα–ligand (Å) | Side chain–ligand (Å) | Main chain–ligand (Å) | Polar contact (≤3.2 Å) | pLDDT (Cα) |
|---|---|---|---|---|---|
| **Glu264** | 7.0 | 2.9 (OE2–O) | 7.0 (CA–O) | Side chain | 98.4 |
| **Glu358** | 7.4 | 2.9 (OE1–O) | 7.4 (CA–O) | Side chain | 98.6 |
| Tyr335 | 4.8 | 3.2 (OH–C) | 4.2 (O–O) | No | 97.5 |
| Thr360 | 4.5 | 3.6 (CG2–O) | 2.9 (O–O) | Main chain | 95.8 |
| Trp395 | 6.6 | 3.3 (NE1–O) | 6.6 (CA–O) | No | 98.0 |
| Asp156 | 6.7 | 3.6 (OD2–C) | 6.5 (N–O) | No | 96.8 |
| Val361 | 3.7 | 4.8 (CG1–O) | 3.7 (CA–O) | No | 92.7 |
| Val362 | 4.6 | 3.6 (CG1–O) | 3.8 (N–O) | No | 89.8 |
| Tyr272 | 8.7 | 3.0 (OH–O) | 8.6 (O–O) | Side chain | 95.4 |
| Cys418 | 5.6 | 3.8 (SG–O) | 5.6 (CA–O) | No | 94.6 |
| Val267 | 6.0 | 3.8 (CG2–O) | 6.0 (CA–O) | No | 94.6 |
| Phe157 | 7.8 | 4.0 (CE2–O) | 6.9 (N–O) | No | 97.6 |
| Cys415 | 5.3 | 4.1 (SG–C) | 5.3 (CA–C) | No | 92.4 |

##### Pose 3

| Residue | Cα–ligand (Å) | Side chain–ligand (Å) | Main chain–ligand (Å) | Polar contact (≤3.2 Å) | pLDDT (Cα) |
|---|---|---|---|---|---|
| **Glu264** | 6.9 | 2.9 (OE2–O) | 6.9 (CA–O) | Side chain | 98.4 |
| **Glu358** | 7.5 | 2.9 (OE1–O) | 7.5 (CA–O) | Side chain | 98.6 |
| Tyr335 | 4.9 | 3.2 (OH–C) | 4.3 (O–O) | No | 97.5 |
| Thr360 | 5.5 | 3.5 (CG2–O) | 3.7 (O–C) | No | 95.8 |
| Trp395 | 6.5 | 3.1 (NE1–O) | 6.5 (CA–O) | Side chain | 98.0 |
| Asp156 | 6.6 | 3.6 (OD2–O) | 6.4 (N–O) | No | 96.8 |
| Val361 | 3.5 | 4.4 (CB–O) | 3.5 (CA–O) | No | 92.7 |
| Val362 | 4.1 | 3.5 (CG2–O) | 3.1 (N–O) | Main chain | 89.8 |
| Tyr272 | 8.6 | 2.9 (OH–O) | 8.5 (O–O) | Side chain | 95.4 |
| Val267 | 6.0 | 3.8 (CG2–O) | 6.0 (CA–O) | No | 94.6 |
| Phe157 | 7.7 | 4.1 (CE2–O) | 6.9 (N–O) | No | 97.6 |
| Cys418 | 6.5 | 4.1 (SG–O) | 6.5 (CA–O) | No | 94.6 |

Distances (Å) are the minimum between heavy atoms of the indicated part of each residue and the ligand in each of the three poses (poses 1–3) obtained with VINA docking; the atom pair is given in parentheses. For each pose, all residues with at least one heavy atom within 4.5 Å of the ligand are listed. Residues in blue are within 4.5 Å in all three poses; the remaining residues are listed below them and appear only in some poses. A polar contact was defined as an N/O pair ≤3.2 Å. Catalytic residues (Glu264 and Glu358) are in bold. pLDDT is the confidence of the AlphaFold model at the Cα atom of each residue. The centroid of the ligand in poses 2 and 3 was displaced by 1.2 and 1.1 Å relative to pose 1 (heavy-atom RMSD 2.9 and 3.5 Å).

#### Table S5: Structural comparison of Glu1, BT3312, MoGlu16 and Xyn30B

##### Structures used:

**BT3312**: crystal structure 5NGL (chain A, GH30-residues 82–427)
**Xyn30B**: crystal structure 6IUJ (chain A, GH30-residues 37–376)
**Glu1**: A0A0D1E2V9v6 AlphaFold model (GH30 residues 100–420)
**MoGlu16**: AF-G4MNS0-F1v6 AlphaFold model (GH30-MGG_14602, residues 76–407)

##### Method used:

**Structural comparison.** All comparisons were based on Cα atoms. Structures were superimposed in PyMOL with two complementary methods. ***Align*** first pairs residues by sequence, using a substitution matrix, and superimposes the structures on those pairs. It then refines the superposition iteratively: at each cycle, the residue pairs that deviate most (by more than 2.0 times the current RMSD) are discarded and the structures are superimposed again, for up to 10 cycles or until no more pairs are discarded. The RMSD reported is that of the pairs retained, which describes the best-fitting core of the domain; it is therefore always given together with the number of residues retained out of the total in Glu1. As the initial pairing depends on sequence, which is poorly conserved between some of the proteins (about 15% identity between Glu1 and Xyn30B), a second, sequence-independent method was used. ***Cealign*** (combinatorial extension) pairs residues from the structures alone: it compares the pattern of Cα–Cα distances within short fragments of each protein and joins compatible fragments into the longest continuous alignment, allowing gaps. It discards no residues, so it aligns more of each domain and gives a higher RMSD than align. The two measures answer different questions: align shows how closely the best-matching core superimposes, and cealign how much of the domain shares the same backbone fold.

##### 1. Global comparison

**Table 1. GH30 domains aligned on Glu1**

| **GH30-Glu1 (residues 100–420, 321 residues)** | | | | | |
|---|---|---|---|---|---|
| **Compared with Glu1** | **Subfamily** | **align, RMSD (Å)** | **align, residues** | **cealign, RMSD (Å)** | **cealign, residues** |
| BT3312 (5NGL, 82–427) | GH30_3 | 1.07 | 236/321 (74%) | 2.76 | 296/321 (92%) |
| MoGlu16 (AlphaFold, 76–407) | GH30_3 | 1.03 | 202/321 (63%) | 2.47 | 288/321 (90%) |
| Xyn30B (6IUJ, 37–376) | GH30_7 | 1.46 | 110/321 (34%) | 3.56 | 280/321 (87%) |
| **GH30-Glu1 (residues 100–540, 441 residues)** | | | | | |
| **Compared with Glu1** | **Subfamily** | **align, RMSD (Å)** | **align, residues** | **cealign, RMSD (Å)** | **cealign, residues** |
| BT3312 (5NGL, 82–427) | GH30_3 | 1.07 | 242/441 (55%) | 2.74 | 304/441 (69%) |
| MoGlu16 (AlphaFold, 76–407) | GH30_3 | 1.03 | 242/441 (55%) | 2.64 | 304/441 (69%) |
| Xyn30B (6IUJ, 37–376) | GH30_7 | 1.46 | 110/441 (25%) | 3.53 | 288/441 (65%) |

***Table 1.*** *Structural similarity of the GH30 domain of each protein to that of Glu1, using two definitions of the Glu1 domain: residues 100–420 (321 residues) and residues 100–540 (441 residues), the latter including the C-terminal α8 helix. For each comparison, the RMSD (Å) of the superimposed Cα atoms is given, together with the number of residues superimposed out of the 321 of Glu1.* ***align*** *(PyMOL; up to 10 refinement cycles, outlier rejection cutoff of 2.0 × RMSD) superimposes the structures and, at each cycle, discards the residue pairs that fit worst.*

**Table 2. Full-length proteins aligned on Glu1 (664 residues)**

| **Compared with Glu1** | **Subfamily** | **align, RMSD (Å)** | **align, residues** | **cealign, RMSD (Å)** | **cealign, residues** |
|---|---|---|---|---|---|
| BT3312 (5NGL) | GH30_3 | 1.28 | 284/664 (43%) | 2.84 | 344/664 (52%) |
| MoGlu16 (AlphaFold) | GH30_3 | 1.04 | 207/664 (31%) | 3.47 | 344/664 (52%) |
| Xyn30B (6IUJ) | GH30_7 | 1.52 | 142/664 (21%) | 3.50 | 304/664 (46%) |

***Table 2:*** *As in Table 1, with the full-length proteins (Glu1, 664 residues). Only the GH30 domain of Glu1 has a counterpart in the other proteins, so a smaller fraction of Glu1 can be superimposed and the cealign RMSD is higher. For Xyn30B, 10 cycles of align were not enough (1.68 Å over 150 residues); the value shown (1.52 Å over 142 residues) is the converged one, which needed 13 cycles.*

**Table 3. Confidence of the Glu1 and MoGlu16 models (pLDDT)**

| **Domain** | **Residues** | **Mean pLDDT** | **Minimum** | **≥ 90** | **≥ 70** |
|---|---|---|---|---|---|
| Glu1 (100–420) | 321 | 96 | 76 | 290 (90%) | 321 (100%) |
| MoGlu16 (76–407) | 332 | 97 | 78 | 318 (96%) | 332 (100%) |

*Table 3: Confidence of the AlphaFold models over the domains used in the comparison: mean and minimum pLDDT, and number of residues with pLDDT ≥ 90 (very high confidence) and ≥ 70 (confident). No residue of either domain is below 70, so the comparison does not rely on poorly predicted regions. The crystal structures have no pLDDT, cristal resolution is 1.85 Å for 5NGL and 2.25 Å for 6IUJ.*

#### 2. Secondary structure and loops

##### Method used:

Secondary structure was assigned from the three-dimensional structures (**PyMOL dss**, applied to the crystal structures of BT3312 and Xyn30B and to the AlphaFold models of Glu1 and MoGlu16), which does not depend on sequence and treats the four proteins in the same way. Helices and strands were then numbered in the canonical order of a TIM barrel (α1–α8 and β1–β8), using the two catalytic glutamates as anchors, since in clan GH-A the acid/base lies at the end of β4 and the nucleophile at the end of β7; equivalent elements share a row in the tables. As dss does not detect every strand, β5 of BT3312 and MoGlu16 was assigned from the **Cα geometry** and from its packing between β4 and β6. unit, the same in the four proteins) and inspected visually, which confirmed that each range corresponds to the intended helix or strand of the three-dimensional structure

**Table 4. α-Helices (canonical TIM-barrel numbering)**

| Helix | BT3312 | Glu1 | MoGlu16 | Xyn30B |
|---|---|---|---|---|
| α1a | 90–98 | 106–118 | 88–96 | 46–51 |
| α1b | 101–112 | 119–131 | 99–110 | 58–69 |
| α2 | 152–168 | 187–197 | 155–170 | 121–131 |
| α3 | 209–226 | 233–252 | 207–224 | 173–190 |
| α4 | 252–270 | 279–296 | 251–269 | 216–233 |
| α5 | 294–307 | 313–322 | 285–298 | 247–258, 267–270 |
| α6 | 320–329 | 339–348 | 313–322 | Absent |
| α7 | 351–369 | 370–386 | 344–354 | 319–333 |
| α8 | 414–425 | 528-440 | 401–412 | 365–376 |

*Table 4: α-Helices of the catalytic domain in the four proteins; equivalent helices are sharing the same row. Secondary structure was assigned from the three-dimensional structures (**PyMOL dss**, applied to the crystal structures and to the AlphaFold models) and the helices were numbered in canonical TIM-barrel order, with the catalytic glutamates (at the end of β4 and β7) as anchors. α1 is a helix–turn–helix (α1a and α1b) in the four proteins. α6 is absent in Xyn30B. The assigned residue ranges were then colored on the structures and inspected visually, which confirmed that each range corresponds to the intended helix or strand of the three-dimensional structure.*

**Table 5. β-Strands**

| Strand | BT3312 | Glu1 | MoGlu16 | Xyn30B |
|---|---|---|---|---|
| β1 | 83–87 | 100–103 | 84–86 | 38–41 |
| β2 | 122–126 | 147–151 | 119–123 | 79–83 |
| β3 | 173–177 | 202–207 | 175–180 | 135–140 |
| β4 **(acid/base)** | 232–234 | 258–260 | 230–232 | 196–198 |
| β5 | 276–281* | 302–305 | 273-280 * | 238–241 |
| β6 | 310–314 | 329–333 | 301–305 | 273–276 |
| β7 **(nucleophile)** | 334–337 | 353–356 | 327–330 | 293–295 |
| β8 | 373–376 | 391–394 | 359–362 | 337–340 |

*Table 5: β-Strands of the barrel; equivalent strands share a row. They were assigned and numbered in the same way as the helices. dss does not detect β5 in BT3312 and MoGlu16, so these two were assigned from the Cα geometry (Cα(i)–Cα(i+2) distances of 6.4–7.3 Å) and from the packing of the segment between β4 and β6.*

#### 3. BT3312 compared with Glu1 and MoGlu16

##### Method used:

**Residue-level comparison with BT3312.** The residues of the BT3312 active site described by Temple et al. (2017) were taken as the reference: the catalytic glutamates (Glu238 and Glu339), the residues of subsite −1 and −2, the positive-subsite residues, the two residues that distinguish BT3312 from other GH30 subfamilies (Asp131 and Asn244), the Loop 2 disulfide (Cys393–Cys396) and the two loops that close the cleft (Loop 1, Gln236–Trp252; Loop 2, residues 381–398). After superposing the structures on Glu1, the equivalent of each residue in Glu1 and MoGlu16 was defined as its reciprocal Cα nearest neighbour within 3.5 Å, and the distance was recorded as Cα–Cα. Only Cα atoms were used, as AlphaFold models, side-chain positions are less reliable. When the equivalent was not of the same type, or when there was none, a candidate was proposed: the closest residue of the same chemical type within

6 Å, graded as reasonable, possible or weak according to chemical similarity and distance. Disulfide bonds were checked from the Sγ–Sγ distance within each protein. The activity of the Ala variants of BT3312 is taken from Temple et al. (2017); none of the candidates has been tested experimentally.

#### Table 6. Positions of BT3312 that are conserved in Glu1 and MoGlu16

| **BT3312** | **Glu1** | **BT3312–Glu1 (Å)** | **MoGlu16** | **BT3312–MoGlu16 (Å)** | **Role in BT3312 (Temple et al., 2017)** |
|---|---|---|---|---|---|
| **Glu238** | Glu264 | 0.4 | Glu236 | 0.4 | Catalytic acid/base; E238A inactive |
| **Glu339** | Glu358 | 0.2 | Glu332 | 0.3 | Catalytic nucleophile; E339A and E339Q inactive |
| **Asp131** | Asp156 | 0.6 | Asp128 | 1.1 | Diagnostic residue, not conserved in GH30_1 or GH30_8 |
| **Trp179** | Trp208 | 0.6 | Trp181 | 0.5 | Subsite −1 (contact with O3) |
| **Asn237** | Asn263 | 0.4 | Asn235 | 0.5 | Subsite −1 (contact with O2) |
| **Trp377** | Trp395 | 0.1 | Trp363 | 0.7 | Subsite −1, hydrophobic platform |
| **Cys393** | Cys415 | 1.7 | Cys381 | 1.0 | Loop 2 disulfide; C393S leaves 0.5% activity |
| **Cys396** | Cys418 | 1.2 | Cys384 | 1.3 | Loop 2 disulfide; C396S leaves 3% activity |
| **Asn282** | Asn309 | 0.6 | Asn281 | 0.7 | Positive subsite; N282A leaves 0.9% activity |

***Table 6.1:*** *Residues of the BT3312 active site described by Temple et al. (2017) that are identical at the equivalent positions of Glu1 and MoGlu16. The equivalent of each residue is its reciprocal Cα nearest neighbour within 3.5 Å after superposition on Glu1, and the distance given is Cα–Cα (Å). The last column gives the role of each residue in BT3312 and, where it was tested, the activity retained by its variant relative to the wild-type enzyme (Cys⟶Ser variants for the disulfide).*

| **Loop 2 disulfide of BT3312 and its equivalents** | | | | | | | |
|---|---|---|---|---|---|---|---|
| **BT3312** | **Glu1** | **BT3312–Glu1 (Å)** | **MoGlu16** | **BT3312–MoGlu16 (Å)** | **Xyn30B** | **BT3312–Xyn30B (Å)** | **Role in BT3312 (Temple et al., 2017)** |
| Cys393 | Cys415 | 1.7 | Cys381 | 1.0 | None (Leu301*) | 7.8* | Loop 2 disulfide; C393S leaves 0.5% activity |
| Cys396 | Cys418 | 1.2 | Cys384 | 1.3 | None (Ser351*) | 3.9* | Loop 2 disulfide; C396S leaves 3% activity |
| Sγ–Sγ 2.12 Å | Sγ–Sγ 2.02 Å | — | Sγ–Sγ 2.01 Å | — | No disulfide | — | |

***Table 6.2:*** *Sγ–Sγ correspond distance between the two sulfur atoms of a Cys pair within one protein (2.0–2.1 Å in a typical disulfide); the rest of the distances corresponde Cα–Cα between equivalent positions after superposition on Glu1. The Loop 2 disulfide of BT3312 is also present in Glu1 and MoGlu16 but not in Xyn30B, and that of Xyn30B is absent from the other three. Green: identical residue. *: nearest neighbour that is not reciprocal or lies beyond 3.5 Å. Sγ–Sγ values come from the crystal structures (BT3312, Xyn30B) and the AlphaFold models (Glu1, MoGlu16).*

#### Table 7. Positions of BT3312 not conserved in Glu1 and MoGlu16, with the nearest candidate

| **BT3312** | **Glu1 equivalent** | **Glu1 candidate** | **MoGlu16 equivalent** | **MoGlu16 candidate** | **Role in BT3312 (Temple et al., 2017)** |
|---|---|---|---|---|---|
| His281 | Phe308 (0.6 Å) Different residue | His333 (5.2 Å) Reasonable candidate | His280 (0.6 Å) | - | Positive subsite; H281A leaves 0.6% activity |
| Ser245 | Tyr272 (0.8 Å) Different residue | - | Met244 (1.1 Å) Different residue | - | Next to Asn244 |
| Asn244 | None | Gln271 (4.3 Å) Possible candidate | None | Asn241 (4.3 Å) Reasonable candidate | Diagnostic residue, not conserved in GH30_1 or GH30_8 |
| Trp345 | None | - | None | - | Subsite −2; W345A leaves 0.4% activity |
| Asp286 | None | Asp311 (5.7 Å) Reasonable candidate | None | - | Positive subsite; D286A leaves 27% activity |

***Table 7****. Residues of the BT3312 active site whose equivalent in Glu1 or MoGlu16 is a different residue or is missing. The equivalent is the residue at the equivalent position after superposition on Glu1 (reciprocal Cα neighbour within 3.5 Å), with its Cα–Cα distance in parentheses; it is shown in green when identical to the BT3312 residue, in orange when conservative, in red when different. When the equivalent is not of the same type, or there is none, a candidate is given: the closest residue of the same chemical type within 6 Å (Cα–Cα distance in parentheses), graded as reasonable (green), possible (orange) or weak (red) according to chemical similarity and distance.*

#### Table 8. Loops 1 and 2 of BT3312 and their equivalents in Glu1 and MoGlu16

| **Loop 1 of BT3312 (Gln236–Trp252)** | | | | |
|---|---|---|---|---|
| **BT3312** | **Glu1** | **BT3312–Glu1 (Å)** | **MoGlu16** | **BT3312–MoGlu16 (Å)** |
| Gln236 | Ala262 | 0.4 | Gln234 | 0.5 |
| Asn237 | Asn263 | 0.4 | Asn235 | 0.5 |

| | | | | |
|---|---|---|---|---|
| Glu238 | Glu264 | 0.4 | Glu236 | 0.4 |
| Pro239 | Pro265 | 0.7 | Pro237 | 0.5 |
| Leu240 | Gln266 | 0.6 | Leu238 | 0.5 |
| Asn241 | Val267 | 0.8 | Asn239 | 0.7 |
| Arg242 | Ala269 | 2.6 | Ser240 | 0.9 |
| Gly243 | Val270 | 1.4 | Asn241 | 0.8 |
| Asn244 | Gln271* | 4.3 | Asn241* | 4.3 |
| Ser245 | Tyr272 | 0.8 | Met244 | 1.1 |
| Ala246 | Pro273 | 1.0 | Pro245 | 0.8 |
| Ser247 | Ser274 | 0.4 | Ser246 | 0.5 |
| Leu248 | Met275 | 0.3 | Leu247 | 0.6 |
| Tyr249 | His276 | 0.6 | Tyr248 | 0.6 |
| Met250 | Leu277 | 0.7 | Val249 | 0.4 |
| Glu251 | Ser278 | 1.8 | Phe250 | 0.8 |
| Trp252 | Ala279 | 1.4 | Pro251 | 0.7 |
| **Loop 2 of BT3312 (residues 381–398)** | | | | |
| **BT3312** | **Glu1** | **BT3312–Glu1 (Å)** | **MoGlu16** | **BT3312–MoGlu16 (Å)** |
| Leu381 | Leu399 | 1.3 | Thr367 | 1.3 |
| Asp382 | Arg400 | 1.5 | Asp368 | 2.2 |
| Asn383 | Lys401 | 1.5 | Asn369* | 2.8* |
| Asp384 | Phe406 | 2.4 | Gln370 | 1.9 |
| Arg385 | Thr407 | 1.2 | Asp371 | 1.3 |
| Gly386 | Thr408 | 1.2 | Gly372 | 0.9 |
| Pro387 | Pro409 | 0.9 | Pro373 | 1.1 |
| Asn388 | His410 | 1.1 | His374 | 1.2 |
| Arg389 | Leu411 | 0.8 | Leu375 | 1.0 |
| Glu390 | Pro412 | 0.8 | Pro376 | 1.7 |
| Gly391 | Lys413 | 2.8 | Gly243* | 3.3* |
| Gly392 | Val414 | 2.1 | Ser380 | 1.2 |
| Cys393 | Cys415 | 1.7 | Cys381 | 1.0 |
| Gln394 | Glu416 | 1.8 | Ala382 | 0.9 |
| Thr395 | Asn417 | 3.1 | Thr383 | 2.2 |
| Cys396 | Cys418 | 1.2 | Cys384 | 1.3 |
| Tyr397 | Asn419 | 2.2 | Thr385 | 1.7 |
| Gly398 | Ala420 | 1.7 | Gly386 | 1.4 |

***Table 8:*** *Loops 1 and 2 of BT3312 and their equivalents in Glu1 and MoGlu16. Residue-by-residue comparison of the two loops that close the active-site cleft of BT3312 (Loop 1, Gln236–Trp252; Loop 2, residues 381–398, stabilised by the Cys393–Cys396 disulfide; Temple et al., 2017). The equivalent of each residue is its reciprocal Cα nearest neighbour within 3.5 Å after superposition on Glu1, and the distance given is Cα–Cα (Å); identical residues are shown in green. For residues with no equivalent (*), the candidate is given instead, i.e. the closest residue of the same chemical type within 6 Å.*

### 4. Xyn30B compared with Glu1 and MoGlu16

##### Table 9. Positions of Xyn30B conserved in several GH30 families

| Xyn30B | Glu1 | Xyn30B–Glu1 (Å) | MoGlu16 | Xyn30B–MoGlu16 (Å) | Role in Xyn30B (Nakamichi et al., 2019) |
|---|---|---|---|---|---|
| Glu202 | Glu264 | 1.0 | Glu236 | 0.4 | Catalytic acid/base (putative; not mutated) |
| Glu297 | Glu358 | 0.2 | Glu332 | 0.2 | Catalytic nucleophile (putative; not mutated) |
| Trp141 | Trp208 | 0.6 | Trp181 | 0.8 | Subsites −1/+1; shared with GH30_8 and with BT3312, so it does not discriminate |
| Asn201 | Asn263 | 0.8 | Asn235 | 0.3 | Subsites −1/+1; shared with GH30_8 and with BT3312 |
| Tyr279 | Tyr335 | 0.1 | Tyr307 | 0.9 | Subsites −1/+1; shared with GH30_8 and with BT3312 |
| Trp341 | Trp395 | 0.7 | Trp363 | 0.7 | Subsites −1/+1 and −2b |

***Table 9:*** *Positions of Xyn30B (catalytic residues and subsite −1/+1) that are conserved in Glu1 and MoGlu16. They are also conserved across other GH30 subfamilies (Temple et al., 2017; Nakamichi et al., 2019), so they do not discriminate between glucanases and xylanases. Distances are Cα–Cα in Å after superposition on Glu1.*

##### Table 10. Other positions and features of Xyn30B compared with Glu1 and MoGlu16

| Xyn30B | Glu1 equivalent | Glu1 candidate | MoGlu16 equivalent | MoGlu16 candidate | Role in Xyn30B (Nakamichi et al., 2019) |
|---|---|---|---|---|---|
| Tyr209 | Tyr272 (0.5 Å), Identical | - | Met244 (0.7 Å), different residue | - | Subsites −1/+1 |
| Arg46 | Val108 (1.6 Å), Different residue | - | Thr90 (1.6 Å), different residue | - | Subsite −2b, recognises MeGlcA; R46A lowers activity 3.2-fold |
| Leu301 | Val362 (2.6 Å), Conservative residue | - | None | - | Subsite −2b |
| Ile342 | Asn396 (1.0 Å), Different residue | - | Thr364 (1.0 Å), Different residue | - | Subsite −2b |
| Glu345 | Leu399 (0.7 Å), Different residue | - | Thr367 (1.7 Å), Different residue | - | Subsite −2b |
| Thr349 | None | Thr363 (3.8 Å) Reasonable candidate | None | - | Subsite −2b |
| Ser351 | Asn419 (1.6 Å), Polar | - | Thr335 (2.9 Å), Conservative residue | - | Subsite −2b |
| Asn93 | Gly158 (1.7 Å), Different residue | - | Pro131 (1.8 Å), Different residue | - | Tip of the β2–α2 loop; associated with exo activity |
| Cys241 | Ala305 (1.7 Å), Different residue | - | Ala277 (1.4 Å), Different residue | - | Disulfide of fungal GH30_7 |
| Cys242 | Trp306 (1.9 Å), Different residue | - | Tyr278 (1.6 Å), Different residue | - | Disulfide of fungal GH30_7 |

***Table 10:*** *Positions of Xyn30B that define xylanase activity or the fungal GH30_7 subfamily (Nakamichi et al., 2019) and their equivalents in Glu1 and MoGlu16. Equivalent and candidate are defined as in Table 9: green marks an identical residue, orange a conservative one, red a different one, and blue shading means there is no equivalent; a candidate is the closest residue of the same type within 6 Å and is graded reasonable, possible or weak.*

### Table S6: Docking of alternative substrates in Glu1 and of β-gentiobiose in BT3312

Docking was performed with SwissDock (AutoDock Vina, sampling exhaustivity 40) on the AlphaFold model of Glu1 (UniProt A0A0D1E2V9) and on the crystal structure of BT3312 (PDB 5NGL, chain A). The search space was a 20 × 20 × 20 Å box centred on the catalytic site, with its centre at (−3, 8, −3) for Glu1 and (1, 6, 18) for BT3312.

#### Table 1. Glu1: calculated affinity of the docking models for each ligand

| β-1,6-glucanase activity CID PubChem: 441422 Ligand: gentiobiose | | β-1,4-xylanase activity CID PubChem: 439538 Ligand: xylobiose | | β-1,3-glucanase activity CID PubChem: 439306 Ligand: laminarin | | Chitinase activity CID PubChem: 439544 Ligand: N,N'-diacetylchitobiose | |
|---|---|---|---|---|---|---|---|
| **Model** | **Calculated affinity (kcal/mol)** | **Model** | **Calculated affinity (kcal/mol)** | **Model** | **Calculated affinity (kcal/mol)** | **Model** | **Calculated affinity (kcal/mol)** |
| 1 | −6.142 | 1 | −5.681 | 1 | −5.408 | 1 | −5.877 |
| 2 | −6.065 | 2 | −5.531 | 2 | −5.271 | 2 | −5.776 |
| 3 | −5.886 | 3 | −5.503 | 3 | −5.100 | 3 | −5.700 |
| 4 | −5.769 | 4 | −5.367 | 4 | −5.098 | 4 | −5.606 |
| 5 | −5.712 | 5 | −5.343 | 5 | −5.038 | 5 | −5.401 |
| 6 | −5.675 | 6 | −5.275 | 6 | −4.939 | 6 | −5.368 |
| 7 | −5.663 | 7 | −5.275 | 7 | −4.936 | 7 | −5.364 |
| 8 | −5.590 | 8 | −5.165 | 8 | −4.821 | 8 | −5.224 |
| 9 | −5.543 | 9 | −5.149 | 9 | −4.813 | 9 | −5.218 |
| 10 | −5.542 | 10 | −5.138 | 10 | −4.781 | 10 | −5.202 |
| 11 | −5.536 | 11 | −5.124 | 11 | −4.747 | 11 | −5.167 |
| 12 | −5.532 | 12 | −5.100 | 12 | −4.743 | 12 | −5.153 |
| 13 | −5.520 | 13 | −5.038 | 13 | −4.729 | 13 | −5.095 |
| 14 | −5.510 | 14 | −5.032 | 14 | −4.727 | 14 | −5.062 |
| 15 | −5.449 | 15 | −5.018 | 15 | −4.718 | 15 | −5.059 |
| 16 | −5.406 | 16 | −5.014 | 16 | −4.717 | 16 | −4.975 |
| 17 | −5.404 | 17 | −4.861 | 17 | −4.712 | 17 | −4.956 |
| 18 | −5.404 | 18 | −4.777 | 18 | −4.695 | 18 | −4.838 |
| 19 | −5.335 | 19 | −4.757 | 19 | −4.651 | 19 | −4.800 |
| | | 20 | −4.662 | 20 | −4.522 | | |

**Table 1:** *Calculated affinity (kcal/mol) of each model obtained with AutoDock Vina, ranked from the highest to the lowest affinity (more negative values indicate stronger binding).*

#### Table 2. BT3312: calculated affinity of the docking models for β-gentiobiose

| β-1,6-glucanase activity<br>CID PubChem: 441422<br>Ligand: gentiobiose | |
|---|---|
| **Model** | **Calculated affinity (kcal/mol)** |
| 1 | −6.869 |
| 2 | −6.802 |
| 3 | −6.662 |
| 4 | −6.573 |
| 5 | −6.569 |
| 6 | −6.520 |
| 7 | −6.505 |
| 8 | −6.498 |
| 9 | −6.496 |
| 10 | −6.482 |
| 11 | −6.466 |
| 12 | −6.437 |
| 13 | −6.435 |
| 14 | −6.402 |
| 15 | −6.393 |
| 16 | −6.390 |
| 17 | −6.388 |
| 18 | −6.268 |
| 19 | −6.226 |
| 20 | −6.205 |

***Table 2:*** *Calculated affinity (kcal/mol) of the twenty docking models of β-gentiobiose in the BT3312 pocket, ranked from the highest to the lowest affinity. The best model was −6.869 kcal/mol.*

### Table S7: Glu1 and UMAG_04357: GH30 core and secondary structure of the $(\beta/\alpha)_8$ barrel

*PyMOL structural alignment (align; 10 cycles, outlier rejection cutoff of 2.0 Å) of the AlphaFold models of Glu1 (UM) and UMAG_04357. Residue numbering of each protein corresponds to its own sequence. RMSD values (Å) were calculated on the backbone atoms (N, Cα, C, O) retained after outlier rejection; the number of residues (retained atoms/4) is given in parentheses.*

#### Table 1. GH30 core defined for Glu1 and UMAG_04357 and structural alignment between them

| Protein | Length (aa) | Predicted GH30 domain | Core defined in this study | Residues in core | RMSD core vs Glu1 (Å) | RMSD full-length vs Glu1 (Å) |
|---|---|---|---|---|---|---|
| **Glu1 (UM)** | 664 | 100–358 | 100–540 | 441 | – | – |
| **UMAG_04357** | 602 | – | 153–502 | 350 | 0.71 (295) | 0.77 (316) |

*The Glu1 core (residues 100–540) spans from the start of the predicted GH30 domain to the end of α8. The UMAG_04357 core (residues 153–502) was defined as the region equivalent to the Glu1 core after superposition of the full-length models. RMSD values were calculated for the two cores superimposed on each other and for the full-length models; in the latter, only 316 residues (about half of each protein) superimpose within the 2.0 Å cutoff.*

#### Table 2. Equivalent β-strands of the barrel

| Element | Glu1 | UMAG_04357 |
|---|---|---|
| **β1** | 99–103 | 153–156 |
| **β2** | 147–151 | 192–196 |
| **β3** | 202–207 | 260–264 |
| **β4** | 258–260 | 311–313 |
| **β5** | 303–308 | 354–357 |
| **β6** | 329–333 | 381–385 |
| **β7** | 353–356 | 409–411 |
| **β8** | 391–394 | 445–448 |

*Strand ranges are those assigned in the AlphaFold models; the Glu1 range in blue (β5) was not detected automatically and is taken from Table SX. Each residue of a Glu1 strand was matched to the closest Cα of UMAG_04357 after superposition (≤4.0 Å); the UMAG_04357 range gives the matched strand. Mean Cα–Cα distances of the matched residues were 0.4–0.8 Å. β5 (354–357) and β7 (409–411) are shorter in UMAG_04357 than in Glu1 as assigned.*

#### Table 3. Equivalent α-helices of the barrel

| Element | Glu1 | UMAG_04357 |
|---|---|---|
| **α1a** | 106–118 | 159–171 |
| **α1b** | 120–131 | 173–184 |
| **α2** | 187–197 | 240–255 |
| **α3** | 233–252 | 287–305 |
| **α4** | 279–296 | 331–348 |
| **α5** | 315–326 | 365–374 |
| **α6** | 339–348 | 391–400 |
| **α7** | 370–386 | 423–440 |
| **Segment β8→α8** | 395–527 (133) | 449–489 (41) |
| **α8** | 528–540 | 490–501 |

*Helices were matched as described for Table 2; mean Cα–Cα distances were 0.6–2.3 Å. The Glu1 range in blue (α5, 315–326) was not detected automatically and is taken from Table SX. Values in parentheses in the β8→α8 row are the number of residues between β8 and α8. α2 is longer in UMAG_04357 (16 vs 11 residues). UMAG_04357 has two additional short helices (217–223 and 227–231) that correspond to coil in Glu1 (172–176) and are not part of the barrel elements.*

### Table S8. Plasmids and oligonucleotides used in this study.

#### Plasmids

| Number | Plasmid | Cloning method | Number | Primer Name | Primer sequence 5'-3' |
|---|---|---|---|---|---|
| 220 | pJET Δ*Um05223*:NatR | Standard molecular cloning | 786 | CoUm05223-5 | GGGAATGTGACAACGCGAGG |
| | | | 787 | CoUm05223-3 | AATCACAGCGCGTCTCTTCC |
| | | | 788 | Um05223 Ko5-1 | CGATGAAAGGGTGCAGATGG |
| | | | 789 | Um05223 Ko5-2 | CACGGCCTGAGTGGCCACCTGTCACTTTCTTGTCCTTCC |
| | | | 790 | Um05223 Ko3-1 | GTGGCCATCTAGGCCCTCAAGAGAAGAGGTCAAGACG |
| | | | 791 | Um05223 Ko3-2 | GCAAGAACTTGACGGTTGAG |
| | | | 1019 | Um05223-Seq-F2_INT5' | GTGGAATCTGGTCTTGAGGAAGG |
| | | | 1021 | Um05223-Seq-R2_INT3' | ATTCGCGGAAAGGTGCATGG |
| 482 | pBSK Δ*Um04357*::GenR | NEBuilder® HiFi DNA Assembly | 2416 | Ko5'_Umag_04357_Fwd | CGAATTCCTGCAGCCCGGGGCGAAAGTAGGCGCAATGC |
| | | | 2417 | Ko5_Umag_04357_Rev | TACCACGTTCGCCGAGCGAGTTTGTTGC |
| | | | 2418 | HygR_Fwd (Umag_04357) | CTCGCTCGGCGAACGTGGTAACTACCAG |
| | | | 2419 | HygR_Rev (Umag_04357) | TCGCGGCACCTCATGTTTGACAGCTTATC |
| | | | 2420 | Ko3_Umag_04357_Fwd | TCAAACATGAGGTGCCGCGACGAGTCAA |
| | | | 2421 | Ko3_Umag_04357_Rev | CGGCCGCTCTAGAACTAGTGCCCGGTGGAGCGTGCCAA |
| | | | 2422 | Co5_Umag_04357 | TGATCGTGGTCGGGAAAC |
| | | | 2423 | Co3_Umag_04357 | TGATCGTGGTCGGGAAAC |
| | | | 2424 | Int5_ Umag_04357 | GAGCGACAGTCAGGGCGT |
| | | | 2425 | Int3_ Umag_04357 | TGTTCGACTTCATAAAGG |
| | | | 2430 | KO5_Umag_04357_PCR | CGAAAGTAGGCGCAATGC |
| | | | 2431 | KO3_Umag_04357_PCR | CCCGGTGGAGCGTGCCAA |
| 466 | p123:P*glu1:glu1* | NEBuilder® HiFi DNA Assembly | 2183 | P:Glu1_Fwd_NEB | acactatagaactcgagcagAATCACAGCGCGTCTCTTCCCTG |
| | | | 2184 | P:Glu1_Rev_NEB | tgaacgatctgcagccgggcCTAGGCTGGCGCAGGCGC |
| 464 | p123:P*glu1:glu1*(E264A/E358A) | NEBuilder® HiFi DNA Assembly | 2247 | PGlu1-Glu1_fwd | acactatagaactcgagcagAATCACAGCGCGTCTCTTC |
| | | | 2248 | PGlu1-Glu1_rev | attcgctgggCTCATCGCATAGATCTCAATCC |
| | | | 2249 | Glu1 (E264A/E358A)_Fwd | atgcgatgagCCCAGCGAATGCCCCTCA |
| | | | 2250 | Glu1(E264A/E358A)_Rev | tgtgtcaccaCGGTTTTTGCGGTAAAGTAGGTATCC |

| Number | Plasmid | Cloning method | Numbe r | Primer Name | Primer sequence 5'-3' |
|---|---|---|---|---|---|
| | | | 2251 | Glu1 (final)_fwd | gcaaaaaccgTGGTGACACAGTGGTTCAACGAGC |
| | | | 2252 | Glu1 (final)_rev | tgaacgatctgcagccgggcCTAGGCTGGCGCAGGCGC |
| 470 | p123 Pglu1:*glu1*::gfp | Standard molecular cloning | 2333 | Fwd-P:Glu1_PvuII (GFP) | AATCACAGCGCGTCTCTT |
| | | | 2334 | Rev-P:Glu1_NcoI (GFP) | AGCCCATGGAGGCTGGCGCAGGCGCAT |
| 217 | p123 Potef:*glu1*::gfp CbxR | Standard molecular cloning | 983 | Um05223startXmaI | AATcccgggATGAGGTCTATCACCAGCTGG |
| | | | 984 | Um05223stopNotI | TTAgcggccgcCTAGGCTGGCGCAGGCGCATCC |

#### Primers_qPCR

*qPCR*

| Name | Sequence (5' - 3') |
|---|---|
| RT-PPI1-5 | ACATCGTCAAGGCTATCG |
| RT-PPI1-3 | CTTCGGCATTGTTGAGGGTTT |
| Gapdh-F | CTTCGGCATTGTTGAGGGTTT |
| Gapdh-R | TCCTTGGCTGAGGGTCCGTC |

*glu1 transcript levels*

| Name | Sequence (5' - 3') |
|---|---|
| glu1 RT-fwd | GAGACAGGTGTTTTCGAGCCAG |
| glu1 RT-rev | CACAGTGGTAACATTCGCCGT |

**Table S9. *U. maydis* strains used in this study.**

| Strain | Genotype | Resistance | Source |
|---|---|---|---|
| FB1 | *a1 b1* | - | Banuett and Herskowitz 1989 |
| FB2 | *a2 b2* | - | Banuett and Herskowitz 1989 |
| FB1 Δ*glu1* | *a1 b1 Δglu1* | NatR | This work |
| FB2 Δ*glu1* | *a2 b2 Δglu1* | NatR | This work |
| CL13 | *a1 bE1 bW2* | - | Bolker et al., 1995 |
| CL13 Δ*glu1* | *a1 bE1 bW2 Δglu1* | NatR | This work |
| CL13 Δ*glu1* P*glu1:glu1* | *a1 bE1 bW2 Δglu1 ipR*[P*glu1:glu1*]$_{ipS}$ | NatR CbxR | This work |
| CL13 Δ*umag_04357* | *a1 bE1 bW2 Δumag_04357* | HygR | This work |
| CL13 Δ*glu1* + P*glu1:glu1*(E264A/E358A) | *a1 bE1 bW2 Δglu1 ipR*[P*glu1*(E264A/E358A):*glu1*]$_{ipS}$ | NatR CbxR | This work |
| CL13 Δ*glu1* 2xRFP | a1 mfa2 bW2 Δ*glu1* ipR[Potef::RFP]$_{ipS}$ | CbxR | This work |
| CL13 CFP | *a1 mfa2 bW2 ipR*[Potef::RFP]$_{ipS}$ | CbxR | This work |
| SG200 | *a1 mfa2 bW2 bE1* | - | Bolker et al., 1995 |
| SG200 Δ*glu1* | *a1 mfa2 bW2 bE1 Δglu1* | NatR | This work |
| SG200 Potef:*glu1*::gfp | *a1 mfa2 bW2 bE1 ipR*[Potef:*glu1*::gfp]*ipS* | NatR | This work |
| SG200 P*glu1:glu1*::gfp | *a1 mfa2 bW2 bE1 ipR*[Pglu1:*glu1*::gfp]*ipS* | NatR | This work |
